# Modeling gene-environment interactions in Parkinson’s Disease: *Helicobacter pylori* infection of *Pink1^−/−^* mice induces CD8 T cell-dependent motor and cognitive dysfunction

**DOI:** 10.1101/2024.02.25.580545

**Authors:** Alexandra Kazanova, Jacqueline Sung, Nathalia Oliveira, Christina Gavino, Sherilyn Recinto, Hicham Bessaiah, Jessica Pei, Lindsay Burns, Willemein Miller, Morgane Brouillard-Galipeau, Lei Zhu, Lucia Guerra, Moustafa Nouh Elemeery, Adam MacDonald, Joel Lanoix, Pierre Thibault, Heidi McBride, Michel Desjardins, Jo Anne Stratton, Nathalie Labrecque, Samantha Gruenheid

## Abstract

Parkinson’s disease (PD) is a chronic neurodegenerative disorder characterized by progressive loss of motor function. Diagnosis occurs late: after motor symptom development downstream of the irreparable loss of a large proportion of the dopaminergic neurons in the *substantia nigra* of the brain. Understanding PD pathophysiology in its pre-motor prodromal phase is needed for earlier diagnosis and intervention. Genetic risk factors, environmental triggers, and dysregulated immunity have all been implicated in PD development. Here, we demonstrate in a mouse model deficient in the PD-associated gene *Pink*, that infection with the human PD-associated gastric bacterium *Helicobacter pylori* leads to development of motor and cognitive signs resembling prodromal features of PD. This was also associated with proliferation and activation of primary mitochondria-reactive CD8 T cells and infiltration of CD8 T cells into the brain. Development of the motor and cognitive phenotypes in the infected *Pink1^−/−^* mice was abrogated when CD8 T cells were depleted prior to infection. We anticipate that this new model, which integrates genetic PD susceptibility, a PD-relevant environmental trigger, and specific immune changes that are required for symptom development, will be a valuable tool for increasing our understanding of this complex disease.

## Introduction

Parkinson’s Disease (PD) is a chronic neurodegenerative disorder characterized by a progressive loss of motor function, involving bradykinesia, tremors, and rigidity (Tysnes & Storstein, 2017). It is further characterized by the loss of dopaminergic neurons and by the formation of α-synuclein-containing Lewy bodies in the *substantia nigra* of the brain (Braak et al., 2003; Tysnes & Storstein, 2017). Decades before motor dysfunction, patients experience non-specific prodromal symptoms, including constipation, depression, behavioral changes, and an increased frequency of bone fractures (Camacho-Soto et al., 2020; Menza et al., 1993; Postuma & Berg, 2019; Roos et al., 2022). PD diagnosis normally occurs following the development of the characteristic motor symptoms, when up to 70% of dopaminergic neurons in the substantia nigra are already lost (Fearnley & Lees, 1991). Thus, there is a strong interest in understanding the pathophysiology of the prodromal period of PD and how it leads to disease onset, in the hopes of developing biomarkers and tools for earlier diagnosis and intervention.

The exact etiology of PD is unknown, although it is thought to be caused by a complex interaction between genetic risk factors and environmental triggers. Familial PD, with a strong genetic component, accounts for only 5-10% of PD cases (Pang et al., 2019). Exposure to various environmental factors, including pesticides, toxins, a dysregulated microbiome and/or microbial infections has also been associated with PD pathogenesis in both humans and in preclinical models (De Miranda et al., 2022; Hey et al., 2023; Liu et al., 2003; Tysnes & Storstein, 2017). One of the most studied microbes associated with PD in humans is *Helicobacter pylori*. A meta-analysis of more than thirty thousand patients suggests that *H. pylori* is associated with the risk of PD (Shen et al., 2017). This microbe is estimated to colonize the stomach of over 50% of the world’s population causing chronic gastritis in most cases. However, in PD patients it tends to be more prevalent (Dobbs et al., 2000; Fu et al., 2020; McGee et al., 2018; Tan et al., 2015; Wei et al., 2024). This and other emerging genetic, epidemiological, and direct experimental lines of evidence point to a role of gastro-intestinal inflammation in PD. For example, people with inflammatory bowel disease have an increased risk of developing PD later in life (Fu et al., 2020; Houser et al., 2018; Lee et al., 2021; Liu et al., 2003), and numerous alterations in the innate and adaptive immune system have been described in PD patients, both in the brain and the periphery (Devos et al., 2013; Houser et al., 2018; Mogi, Harada, Kondo, et al., 1994; Mogi, Harada, Riederer, et al., 1994). Understanding the interplay of the multiple PD susceptibility factors and various environmental inputs will be key in developing a global understanding of the disease process.

One of the genes associated with early onset familial PD is PTEN-induced kinase 1 (PINK1) (Matsuda et al., 2013). This serine/threonine kinase recruits another PD associated protein, the E3 ubiquitin ligase Parkin, to mitochondria (Kane et al., 2014). These two proteins have been implicated in the removal of damaged mitochondria (mitophagy) leading to the hypothesis that the loss of PINK1 function leads to the accumulation of dysfunctional mitochondria in dopaminergic neurons, resulting in their death (Matsuda et al., 2013; Pickrell & Youle, 2015). Yet, *Pink1*-deficient mice display normal levels of basal mitophagy (McWilliams et al., 2018) and do not experience dopaminergic neuron death at steady state (Gispert et al., 2009) suggesting that additional factors or pathways are likely involved.

We and others have recently reported independent roles for PINK1 in the regulation of the immune system. Matheoud *et al*. demonstrated that in antigen presenting cells (APC), PINK1 represses the presentation of self-mitochondrial antigens on MHC class I molecules, such that loss of PINK1 function leads to the expansion of autoreactive CD8 T cells, both *in vitro* and *in vivo* in response to LPS and the Gram-negative mouse intestinal pathogen *Citrobacter rodentium* (Matheoud et al., 2019; Matheoud et al., 2016). Notably, *Pink1*^−/−^ mice infected with *C. rodentium* early in life later develop a dopamine-dependent PD-like motor phenotype, which correlates with an increase in autoreactive CD8 T cells in the periphery and entry of CD8 cells into the brain. PINK1 has also been proposed to be involved in suppression of the acquired immune response through an effect on regulatory T cells. *In vitro* generated induced regulatory T cells (iTregs) from *Pink1*^−/−^ mice display diminished expression of the characteristic cell surface markers CD25 and CTLA-4, with the same level of FoxP3 expression, and a trend towards impaired suppressor function *in vitro* (Ellis et al., 2013). Together, these studies provide evidence that familial PD involving defects in PINK1 may have an adaptive immune component.

The role of adaptive immunity in sporadic PD is also increasingly recognized (Jiang et al., 2018). Post-mortem studies show a significantly greater density of CD8 T cells (Brochard et al., 2009) and higher levels of pro-inflammatory cytokines in the substantia nigra of PD patients’ brain (Mogi, Harada, Kondo, et al., 1994). A recent study using single nuclei sequencing and spatial transcriptomics revealed increased presence of clonally expanded tissue resident memory CD8 T cells in the substantia nigra of PD patients that had altered interaction with endothelial cells and myeloid cells through TCR - MHCI (Jakubiak et al., 2024). Additionally, the relative abundance of peripheral CD8 T cells to the number of CD4+CD25+ T regulatory cells (Tregs) was found to be significantly higher in PD patients (Baba et al., 2005; Thome et al., 2021). In Tregs of PD patients, the expression of FoxP3 and CD25 were found to be reduced, and this correlated with the activation of proinflammatory T cells, suggesting that Tregs in PD patients have an impaired suppressive function (Thome et al., 2021). It has also been reported that PD patients have significantly higher serum levels of tumor necrotic factor alpha (TNF-α) and a tendency to have higher expression of proinflammatory cytokines (Koziorowski et al., 2012; Thome et al., 2021). Overall, these studies suggest a link between both familial and idiopathic PD and dysregulated adaptive immunity.

Building on our previous finding that *Pink1*^−/−^ mice develop a PD-like motor phenotype following the additional environmental trigger of infection with *C. rodentium*, here we investigate the immune consequences in a model that incorporates both PD genetic predisposition (PINK1 deficiency) and a major PD-associated human-relevant pathogen (*H. pylori*).

## Results

### Absence of PINK1 in bone marrow-derived dendritic cells leads to increased mitochondrial antigen presentation in response to H. pylori

We have shown previously that the process of mitochondrial antigen presentation (MitAP) by dendritic cells is induced by Gram-negative bacteria or its cell wall component – lipopolysaccharide (LPS). Moreover, this process is repressed by the PD-associated proteins PINK1 and Parkin, and therefore amplified in their absence. Considering the implication of the Gram-negative bacteria *Helicobacter pylori* in PD we aimed to test whether this pathogen is capable of inducing MitAP as well the extent of PINK1 involvement in this process. For the detection of MitAP, we used the 2cz hybridoma cell line derived from 2C TCR Tg mice (Sha et al., 1988) (Fig. 1A). Over 90% of CD8 T cells of 2C TCR Tg mice recognize epitopes derived from mitochondrial proteins: the 2-oxoglutarate dehydrogenase (OGDH) peptide LSPFPFDL in complex with MHC class I molecule H2-L^d^, and with much lower affinity in complex with H2-K^b^; and the cytochrome C oxidase subunit NDUFA-4 peptide EQYKFYSV (Dutz et al., 1994; Sykulev et al., 1994; Tallquist et al., 1996). These cells also recognize the artificial peptide SIYRYYGL in complex with H2-K^b^ with the highest affinity (Udaka et al., 1996).

**Figure 1.**
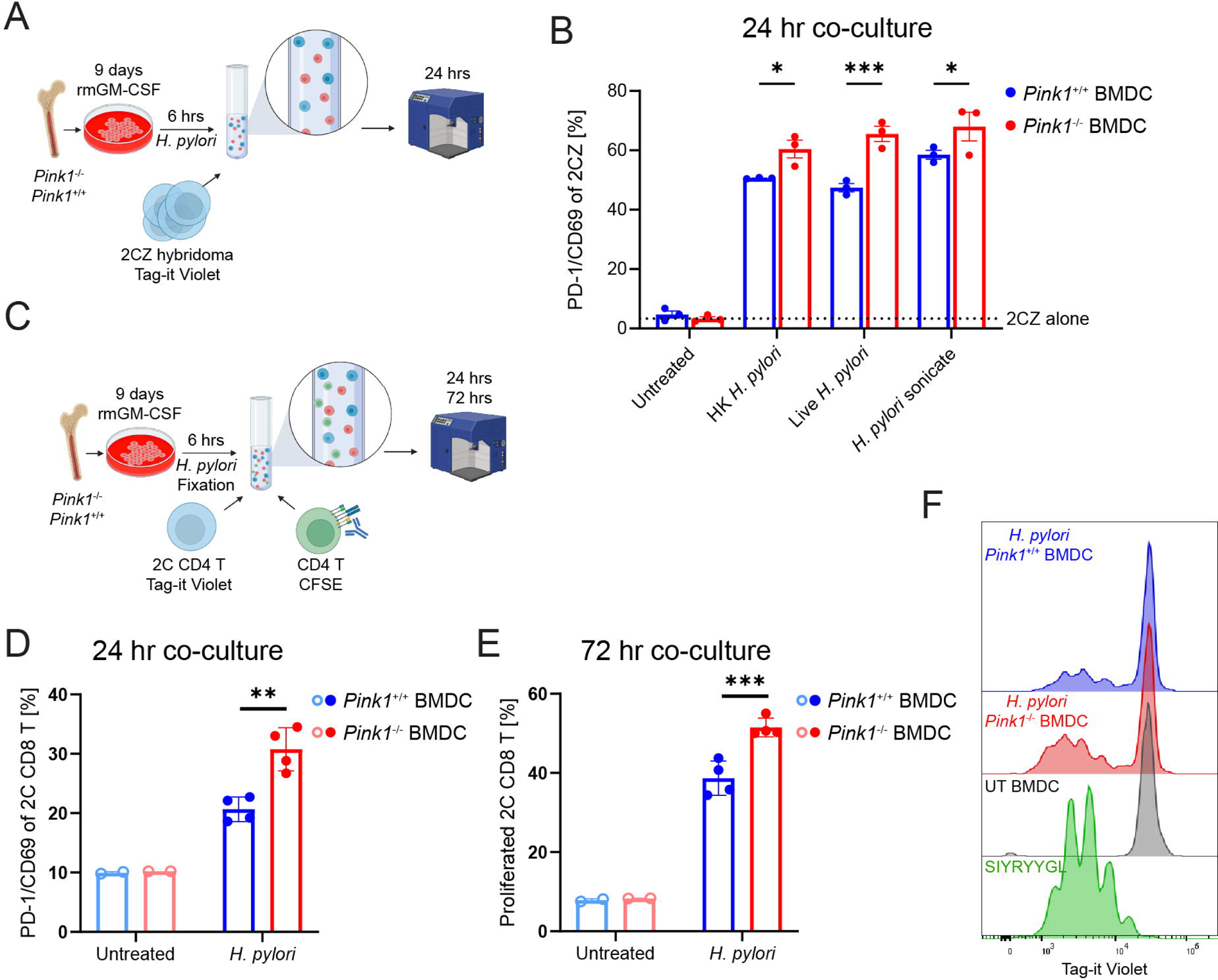
*Pink1*-deficient (*Pink1^−/−^*) bone marrow dendritic cells (BMDCs) have increased mitochondrial antigen presentation (MitAP) in response to *H. pylori*. A. Experimental design. Bone marrow from three *Pink1^−/−^* or WT (*Pink1^+/+^*) littermate mice was isolated and differentiated into BMDCs with 20 ng/ml recombinant mouse GM-CSF (rmGM-CSF) for 9 days. Then, cells were stimulated for 6 hours with either live, heat killed, or sonicated *H. pylori*. For MitAP evaluation, BMDCs were co-cultured with the 2cz hybridoma cell line (derived of 2C TCR Tg mice) for 24 hours and stained for flow cytometry. B. Average frequency of activation-induced marker expression on 2cz cells after 24 h of co-culture. Data are presented as mean ± SEM of technical replicates and analyzed by two-way ANOVA with Sidak post-test. *p<0.05, ***p<0.001. n=3 individual mice per group. The experiment was repeated twice, independently and results from a single representative experiment are shown. C. Experimental design. Bone marrow from three *Pink1^−/−^* or WT (*Pink1^+/+^*) littermate mice was isolated and differentiated into BMDCs with 20 ng/ml recombinant mouse GM-CSF (rmGM-CSF) for 9 days. Then, cells were stimulated for 6 h with sonicated *H. pylori* and fixed. Fixed BMDC were co-cultured with splenic 2C CD8 T cells from two 2C TCR mice labeled with Tag-it Violet cell tracker and splenic CD4+CD25-T cells pooled from three *Pink1* WT mice, preactivated for an hour with anti-CD3 antibody and labeled with CFSE cell tracker. Cells were co-cultured for 24 or 72 hours and stained for flow cytometry. D. Frequency of activation-induced marker expression on 2C CD8 T cells after 24 h of co-culture. E. Frequency of proliferation (cells underwent at least one division) of splenic 2C CD8 T cells after 72 h of co-culture. F. Overlaid histograms showing dilution of Tag-it Violet dye used to assess proliferation of 2C. Data are represented as mean ± SEM and analyzed by two-way ANOVA with Sidak post-test. **p<0.01, ***p<0.001. n=3 individual mice per group. Experiments were done twice, independently.

Previously, quantification of the number of IL-2 producing 2cz hybridoma cells in an ELISPOT assay was used as a readout for MitAP (Matheoud et al., 2016). Here, we adapted the assay for the detection of activation-induced markers (AIM) on 2cz cells with flow cytometry. Indeed, upon presentation of the peptide SIYRYYGL, 2cz cells produced IL-2 and upregulated the AIM-implicated cell surface markers Programmed Cell Death Receptor 1 (PD1), CD69, and CD137 in a dose dependent manner (S.Fig. 1A). Furthermore, IL-2 production in ELISPOT strongly correlated with detection of AIM on 2cz by flow cytometry, thus validating the adaptation of this assay. (S.Fig. 1B).

To test whether *H. pylori* could induce MitAP and whether this was modulated by PINK1, BMDCs from *Pink1****^−/−^*** or WT littermate mice were exposed for six hours to either live, heat killed or sonicated *H. pylori*, and were then co-cultured with 2cz cells. Flow cytometry analysis revealed an upregulation of 2cz AIM following all three stimulations. Moreover, in all *H. pylori*-stimulated conditions, activation of the 2cz cells was significantly higher when the BMDCs were derived from *Pink1****^−/−^*** mice compared to BMDCs from WT mice (Fig.1B).

Activation of naïve T cells *in vivo* depends on three distinct signals: antigen recognition by the T cell receptor; engagement of co-stimulatory molecules on antigen presenting cells with their cognate receptors on T cells; and the presence of specific cytokines that are necessary for T cell proliferation and differentiation. In contrast, activation of the 2cz CD8 hybridoma cell line doesn’t require co-stimulation or cytokine secretion. In order to assess MitAP in an assay that incorporated these signals, we tested whether *H.pylori-*exposed BMDCs could trigger activation of mitochondria-reactive primary CD8 T cells. BMDCs from *Pink1****^−/−^*** or WT littermate mice were exposed to *H. pylori* sonicate for six hours, fixed and co-incubated with primary CD8 T cells isolated from spleens of 2C TCR Tg mice. To provide a source of cytokines, anti-CD3 activated, CD4+CD25-T cells from *Pink1* WT mice were also added to the co-culture. Proliferation of both CD8 and CD4 T cells in the co-cultures was monitored with the use of two distinct cell-tracking dyes.

After exposure to *H. pylori*, BMDCs from both *Pink1****^−/−^*** or WT mice upregulated MHC class I and II molecules as well as the costimulatory molecules CD80 and CD86, but with significant differences in the levels of these molecules between genotypes (S.Fig. 1D). *Pink1****^−/−^*** BMDCs had a distinct signature of lower MHCII, CD80, and CD86 but increased MHCI and PDL1 at the cell surface. Unlike untreated BMDCs which did not stimulate proliferation of the 2C CD8 T cells, BMDCs exposed to sonicated *H.pylori* induced both activation and proliferation of the primary 2C CD8 T cells. Moreover, this process required the presence of activated CD4 T cells (data not shown) and was significantly higher in co-cultures with BMDCs originating from *Pink1****^−/−^*** mice. (Fig. 1C-F). As a control, CD8 T cells from OT-1 TCR transgenic mice, that express a TCR recognizing an unrelated epitope from chicken ovalbumin, did not proliferate in these co-culture conditions (data not shown).

Our results suggest that the stronger induction of MitAP by *H. pylori* sonicate in the absence of PINK1 is mainly attributable to the higher level of MHC-I–self peptide complexes. While MHCI expression was significantly higher in BMDCs from *Pink1****^−/−^*** mice following stimulation, the co-stimulatory molecule expression at the surface of these cells was lower (S.Fig. 1D). 2cz hybridoma activation also strongly correlated with MHCI expression on APCs (S.Fig. 1E). Moreover, the CD4 T cells in the assay, activated independently of BMDC MHC molecules, proliferated independently of BMDC genotype (S.Fig. 1C). Also, the cytokines in the system depended on the CD4 T cells only, as the BMDCs were fixed prior to co-culture. Taken together, these results demonstrate that exposure of antigen presenting cells to the PD-associated pathogen *H. pylori* induced MitAP *in vitro* and that primary autoreactive CD8 T cells can be primed and expanded in response to these cells. While MitAP is induced in response to *H. pylori* in WT antigen presenting cells, this process is more strongly induced when the antigen presenting cells are deficient in the PD-associated protein PINK1.

### Helicobacter pylori infection induces motor-behavioral dysfunction associated with autoreactive CD8 T cells in Pink1−/− mice

To assess the interaction of *H. pylori* exposure with PD genetic susceptibility *in vivo*, we infected *Pink1****^−/−^*** or WT mice with *H. pylori.* Using a previously established qPCR assay to assess colonization (Fox et al., 2007), we showed that *H. pylori* was equally abundant in the stomach of *Pink1****^−/−^*** or WT mice at 2 months post-infection (Fig. 2 A, B). At this time point, *H. pylori* caused only mild gastric pathology (S.Fig. 2A, B, C). No weight loss was observed (data not shown). Although the proinflammatory cytokines IL-17, IL-1β, G-CSF and RANTES were slightly increased in the stomach tissue of infected mice at this time point, we did not detect any significant differences between infected *Pink1 ^−/−^* and WT mice in the level of gastric inflammation (S.Fig. 2D).

**Figure 2.**
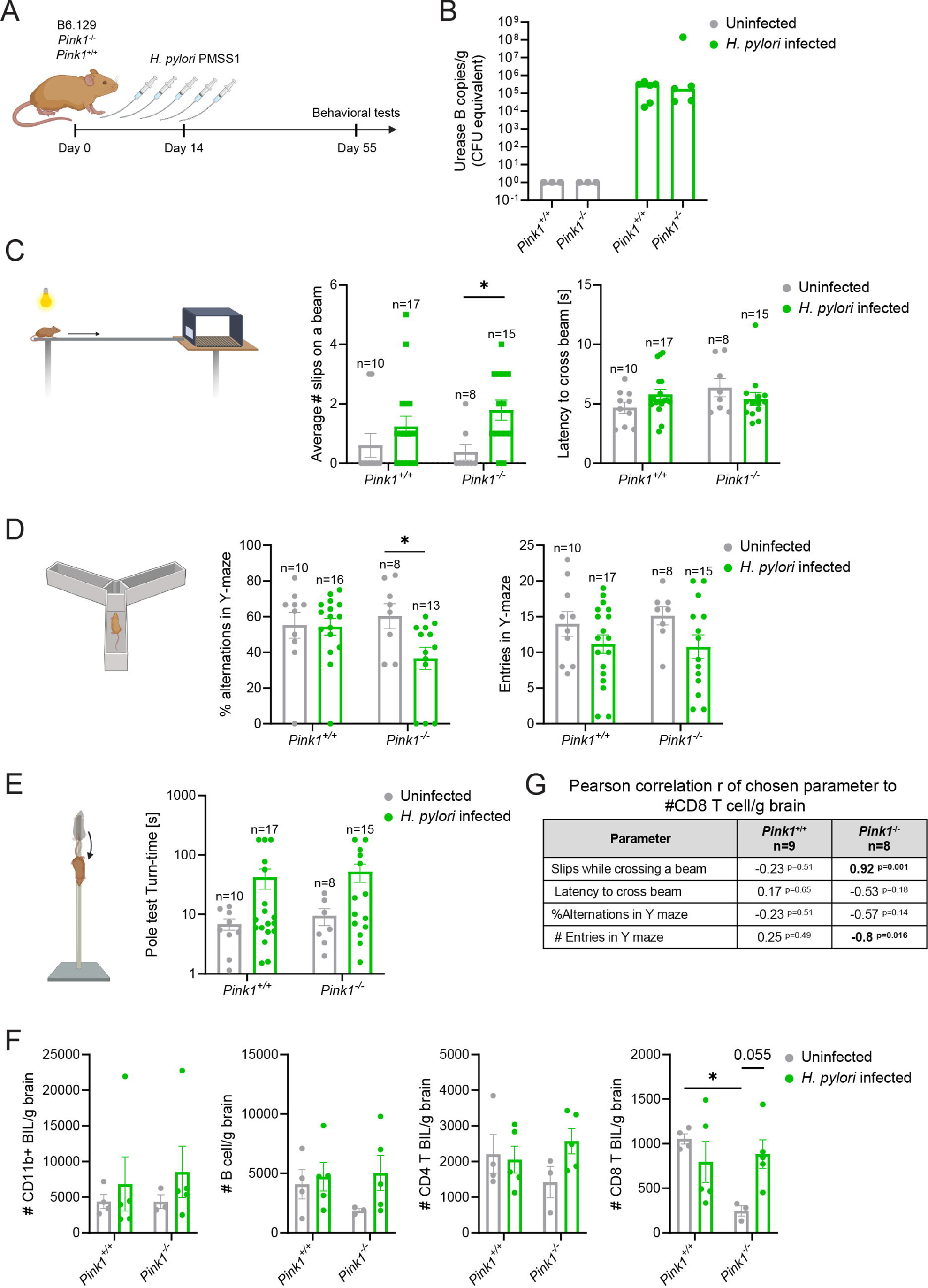
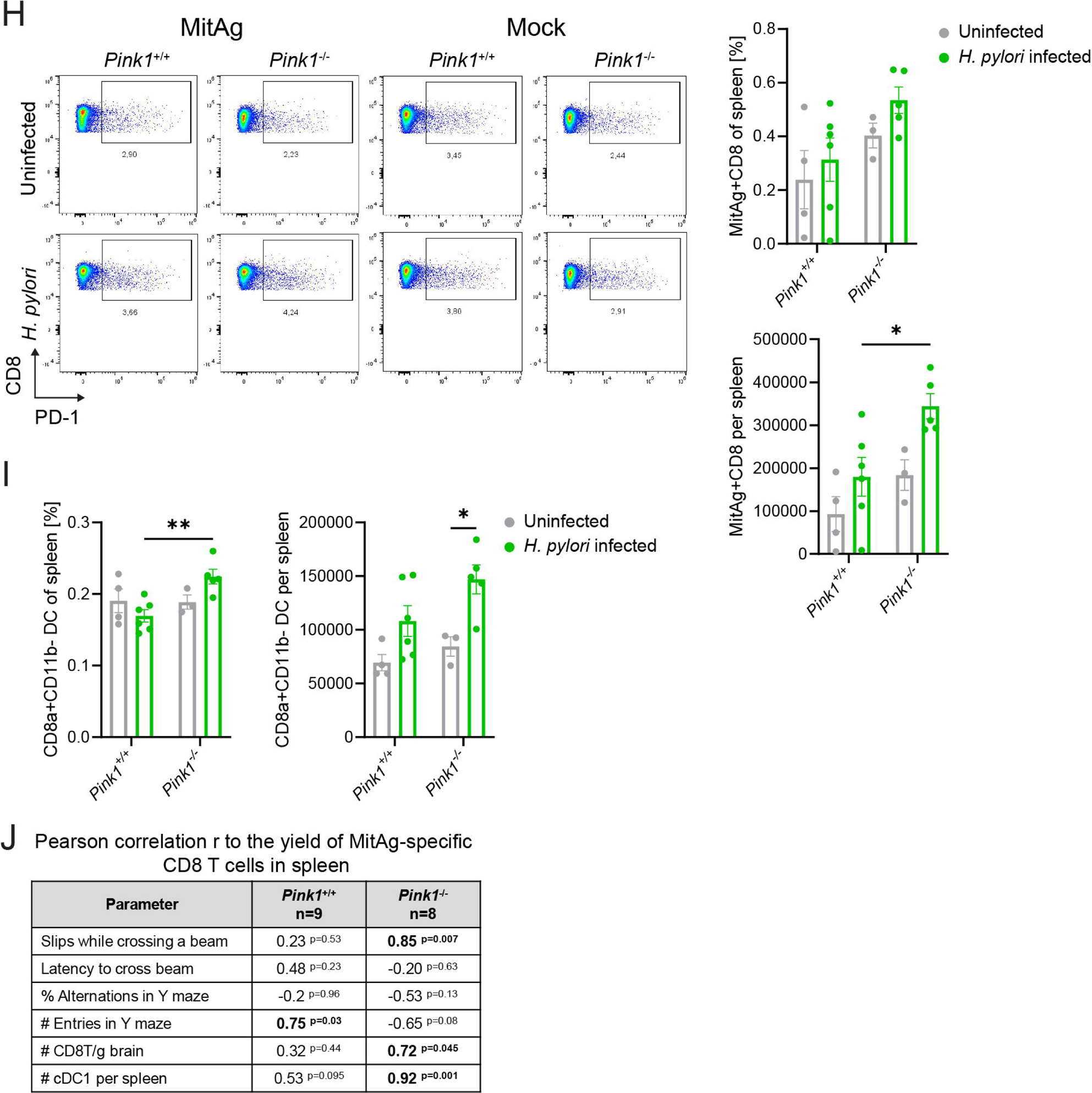
*In vivo H. pylori* infection induces motor-behavioral dysfunction associated with the autoreactive CD8 T in *Pink1^−/−^* mice. A. Experiment design. Six to 10-week-old *Pink1^−/−^* or WT (*Pink1^+/+^*) littermate mice were infected with *H. pylori* PMSS1 strain by oral gavage in a regimen of 5 gavages within 2 weeks. After 2 months of the first gavage, mice were tested in behavioral tests and then sacrificed and had their stomachs, brains, and spleen harvested for further evaluations. B. Average CFU equivalent of *H. pylori* by qPCR of Urease B in stomach homogenates performed in technical triplicates. Data are represented as median and analyzed by two-way ANOVA with Sidak post-test. Infection was performed in mice three individual times with 3-7 mice per group. C-E. Motor and behavioral tests of mice from the two individual representative experiments with at least 4 mice per group at 2 months post infection. C. Beam test – left test overview; middle - average number of slips on a beam; and right - latency to cross the beam; D. Y-maze – left test overview; middle-percentage of alternations; and right - number of entries in each arm. E. Pole test - time to turn on a vertical from upside down to start descending the pole. Data are represented as mean ± SEM and analyzed by two-way ANOVA with Sidak post-test of UI vs *H*.*pylori* condition. Number of mice per group indicated in the figure. *p<0.05. F. Before sacrifice, mice were injected with anti-CD45 in PBS. Brains were collected, weighted and immune cells were isolated, stained for flow cytometry, and brain infiltrating leukocytes were accessed. Data are represented as mean ± SEM and analyzed by two-way ANOVA with Sidak post-test. *p<0.05. n=3-5 mice per group. One representative of three independent experiments is shown. G. Table of Pearson correlation of CD8 T cell brain infiltration and motor test, bold – strong and significant, exact p value is indicated. H. BMDCs were isolated and coated with a self-peptide pool or a mock pool, fixed, and co-cultured with primary splenocytes cells overnight. Then, flow cytometry was performed. Frequency of MitAg+ splenic CD8 T cells and number of MitAg+ CD8 T cells per spleen was determined. H. Upper panel - Down sampled to 5,000 live CD8+ T cell of each mouse in a group concatenated together and represented in a dot plot of AIM (PD-1) expression to the expression of CD8a Left to mitochondrial peptide pool, right to mock pool. Lower left - MitAg reactive CD8 T cell to the total splenocytes increase with *H. pylori* infection, Lower right yield of MitAG+ CD8 t cells per spleen. I. Proportion (left) and yield (right) of cDC1 in spleen Data are represented as mean ± SEM and analyzed by two-way ANOVA with Sidak post-test. 3-5 mice per group. *p<0.05, **p<0.01. J. Table of Pearson correlation (r) of yield of MitAg-specific splenic CD8 T cells to the performance in a motor test, brain CD8 T cell infiltration and cDC1 yield per spleen, bold-p<0.05; exact p values are indicated. One representative of two independent experiments is shown.

Notably, the *H. pylori-*infected *Pink1****^−/−^*** mice developed subtle but significant changes in their motor function and behavior at two months post-infection (Fig. 2C-E). While the latency to cross a beam didn’t change with infection, infected *Pink1****^−/−^*** mice made significantly more slips while crossing the beam than their uninfected counterparts (Fig. 2C). The same tendency was observed in WT mice, but this did not reach the extent nor the statistical significance that was observed in the *Pink1****^−/−^*** mice.

We next performed a Y-maze test to assess cognitive function and anxiety in mice. In infected *Pink1****^−/−^*** mice, the percent of alternations (choice of going to the new area in the maze) was significantly decreased, showing that besides the slips, *H. pylori* infection of *Pink1****^−/−^*** mice causes a cognitive change resulting in decreased willingness to explore new environments (Fig. 2D). The overall number of entries to Y maze arms had a non-significant tendency to be decreased in infected mice of both *Pink1* genotypes.

In our previous work, striking PD-like dopamine-sensitive motor phenotypes developed in *Pink1****^−/−^*** mice six months after a series of gut infections with Gram-negative *Citrobacter rodentium* (Matheoud et al., 2019). This was quantitated using a pole test, where the time it takes a mouse to turn and descend a vertical pole is measured. In contrast, while the *H. pylori-*infected mice of both genotypes tended to have slower descent times at 2 months post-infection, these changes were not significant and no differences between genotypes was observed (Fig. 2E). We hypothesize that at this early time point following infection, we are observing the emergence of subtle signs of dysfunction and that more time would be needed for progression to more pronounced motor phenotypes such as pole test and increases in the time needed to cross a beam.

Having observed these phenotypes in infected *Pink1****^−/−^*** mice, we next investigated if changes in infiltration of immune cells in the brain were also present. After pre-labeling the circulating leukocytes using anti-CD45 injection, we extracted brain infiltrating leukocytes (BIL) by means of gradient centrifugation and analyzed them by flow cytometry. In this experiment, the uninfected *Pink1****^−/−^*** mice had a lower brain infiltration of CD8 T cells compared to WT uninfected mice. (Fig. 2F). However, following *H. pylori* infection, there were no differences in the quantity of BILs in infected *Pink1****^−/−^*** and WT mice. Despite the similar levels of CD8 T cells in the brain following infection between genotypes, the CD8 infiltration strongly and significantly correlated with the motor-behavioral dysfunction in the *Pink1****^−/−^*** mice but not WT mice (Fig. 2G). In the *Pink1****^−/−^*** mice CD8 T cell brain infiltration was strongly associated with slips on the beam (r=0.92; p=0.001) and entries in the Y maze (r=-0.8; p=0.016). The percentage of alternations and the latency to cross the beam did not reach significance. No correlations were observed for the *Pink1* WT mice, leading us to question whether differences in the specificity of the immune cells entering the brain, and particularly the increased presence of anti-mitochondrial CD8 T cells, might be influencing the development of phenotypes in the infected *Pink1****^−/−^*** mice. The yield of CD8 cells infiltrating the brain of mice limited our capacity to do further functional analyses. Accordingly, to test for the emergence of CD8 T cell autoreactivity with *H. pylori* infection, we used restimulation of the splenocytes with a mitochondrial peptide pool (S.Fig 2E). In brief, the BMDCs were coated with a mitochondrial antigen (MitAg) peptide pool and fixed. An irrelevant pool of MHCI H2-K^b^; H2-D^b^ restricted peptides was used as a control. After overnight co-culture with splenocytes from uninfected or infected *Pink1****^−/−^*** or WT mice, the frequency of AIM positive CD8 T cells was determined by flow cytometry. We observed a tendency of an increased proportion, and a significant increase in the total number of MitAg-reactive CD8 T cells in the spleens from infected *Pink1****^−/−^*** mice, compared to uninfected mice or WT infected mice (Fig. 2H). Moreover, the number of cross-presenting classical dendritic cells type I (cDC1) increased in infected mice deficient for PINK1. Correlation analysis showed that the number of auto-reactive CD8 T cells in the spleens of *Pink1****^−/−^*** mice was significantly and strongly associated with motor phenotypes, CD8 T cell brain infiltration, and cDC1. In the case of WT mice, these correlations were not observed (Fig. 2J). This suggested that despite the tendency to generate autoreactive CD8 T cells in WT infected mice there might be some other protective mechanism preventing the development of motor dysfunction. Overall, these results demonstrate that *in vivo H. pylori* infection leads to a systemic increase in autoreactive CD8 T cells, and in the absence of PINK1 these auto reactive CD8 T cells are strongly associated with the global increase in brain-infiltrating CD8 T cells and with a significant motor-behavioral dysfunction.

### H. pylori infection affects regulatory CD4 T cell phenotypes

Knowing the importance of anti-inflammatory CD25+FoxP3+ regulatory T cells (Treg) in the mucosal response to *H. pylori* and the proposed Treg dysfunction in PD, we wanted to investigate whether this population is affected in *Pink1****^−/−^*** mice, and whether *H. pylori* might impact this (Kao et al., 2010; Thome et al., 2021). In both genotypes of mice, *H. pylori* infection led to mild yet significant increase in spleen cellularity (Fig. 3A) showing a systemic effect of the locally mild gastric infection. The frequency of CD25+FoxP3+ T cells among CD45+ leukocytes in the spleen did not differ between genotypes and conditions (Fig. 3B). However, we noted that the intensity of anti-FoxP3 staining in splenic Tregs was significantly decreased in both genotypes following infection (Fig. 3C), whereas the CD25 expression only decreased in *Pink1****^−/−^***-infected mice (Fig. 3D).

**Figure 3.**
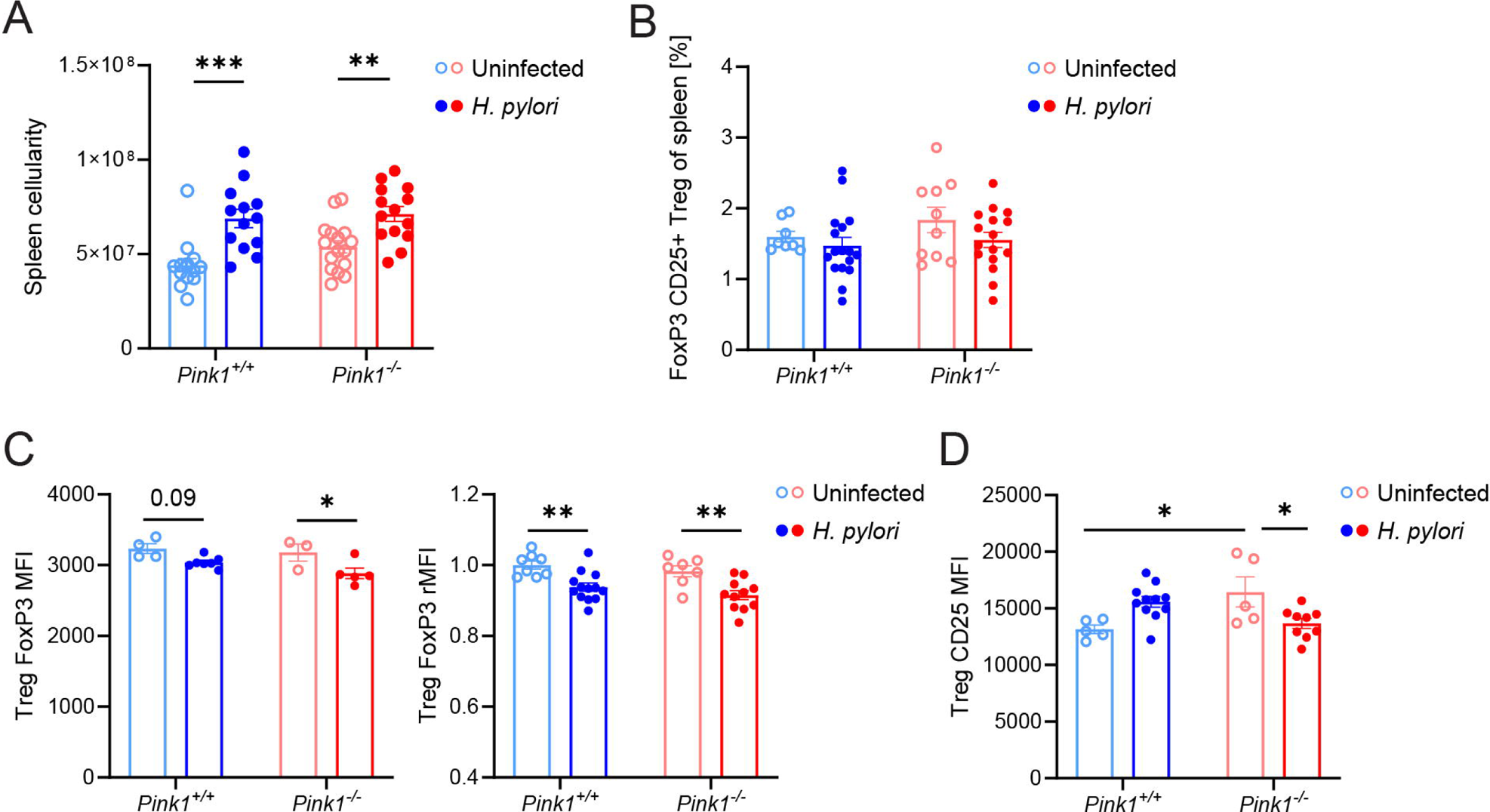
*Pink1^−/−^* mice have altered regulatory T cells (Treg) phenotype after *H. pylori* infection. Six to 10-week-old *Pink1^−/−^* or WT (*Pink1^+/+^*) littermate mice were infected with *H. pylori* PMSS1 strain by oral gavage in a regimen of 5 gavages within 2 weeks. After 2 months of the first gavage, mice were tested in motor behavioral tests and sacrificed and spleens were harvested, and flow cytometry of isolated cells was performed. A. Spleen immune cell count. n=12-16 mice per group. Three independent experiments are represented. B. Frequency of T regs in spleens. n=8-17 mice per group. Two independent experiments are represented. C. MFI of FoxP3 of one individual experiment (left) and relative MFI of FoxP3 in splenic CD25+CD4T cells from two experiments (right) D. CD25 of CD25+FoxP3+ splenic Treg cell at 2mpi, one of two representative individual experiments. n=3-13 mice per group. Data are represented as mean ± SEM and analyzed by two-way ANOVA with Sidak post-test. *p<0.05, **p<0.01, ***p<0.001 or exact p value is shown.

To further investigate the Treg phenotype and its possible modification by *H. pylori*, we exposed splenocytes from uninfected *Pink1****^−/−^*** and WT mice to different concentrations of *H. pylori* sonicate overnight and measured FoxP3 expression. Similar to what was observed *in vivo*, this led to a dose-dependent loss of FoxP3 expression in the Treg population in both genotypes (Fig. 4A). Previously direct effect of *H.pylori* components on human blood CD4 T cells function was also reported (Beigier-Bompadre et al., 2011). Notably, this effect was more pronounced in *Pink1****^−/−^*** Tregs which displayed a significantly higher loss of FoxP3 expression after overnight incubation, even in the untreated condition, suggesting that the stability of FoxP3 might be compromised in the absence of PINK1.

**Figure 4.**
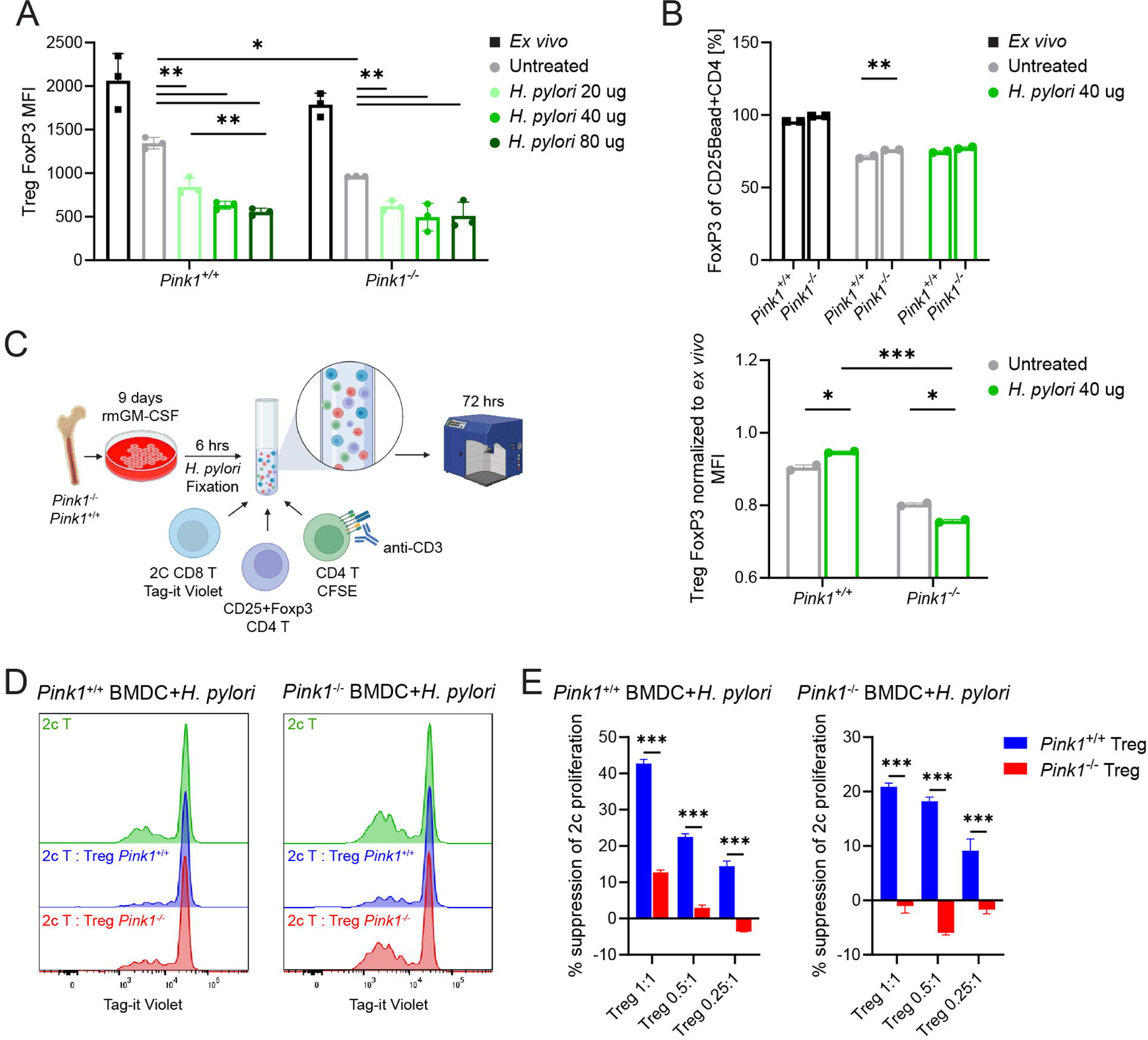
Treg cells of *Pink1^−/−^* mice when exposed to *H. pylori* have dysfunctional phenotype and are defective in suppressor function *in vitro*. *Pink1^−/−^* or WT (*Pink1^+/+^*) littermate mice were sacrificed, and spleens were harvested. Total splenocytes or magnetically isolated CD25+CD4+ Treg population were exposed to sonicated *H. pylori* overnight. Cells were harvested and stained for flow cytometry. A. Dose-response of FoxP3 MFI in Treg cells of *H. pylori*-exposed splenocytes. n=3 mice per group. Experiment performed twice, the representative experiment is shown B. Frequency of FoxP3+ cells (top) and normalized to *ex vivo* overnight level of MFI of FoxP3 of purified Treg cells exposed to *H. pylori* sonicate or left untreated (bottom). Tregs cells were pooled from three *Pink1^−/−^* or three WT (*Pink1^+/+^*) mice and ran in technical duplicates. Data are represented as mean ± SEM and analyzed by two-way ANOVA with Tukey post-test. *p<0.05, **p<0.01, ***p<0.001. One representative of two independent experiments is represented. C. Experimental design. BMDCs from *Pink1^−/−^* or WT (*Pink1^+/+^*) littermate mice (n=3 per group) were isolated and differentiated with 20 ng/ml rmGM-CSF for 9 days. Then, cells were stimulated for 6 hours with sonicated *H. pylori* and fixed. After that, they were co-cultured with splenic 2C CD8 T cells labeled with Tag-it Violet, CFSE-labeled CD4 T cells, previously activated for one hour with soluble anti-CD3 antibody, and different amounts of CD25+CD4 Tregs of three *Pink1^−/−^* or three WT (*Pink1^+/+^*) littermate mice. Incubation occurred for 72 hours, and proliferation and suppression were acquired by flow cytometry. D. Summary histogram of 2C T cells proliferation and suppression by Treg cells. E. Percent of 2C proliferation suppression in the different 2C to Treg ratios. Data represented as mean ± SEM and analyzed by two-way ANOVA with Tukey post-test. *p<0.05, **p<0.01, ***p<0.001. One representative of three independent experiments is shown.

To rule out the potential Treg extrinsic effect of *H pylori* on FoxP3 of Tregs, we magnetically isolated CD25+CD4 T cells from the spleens of the uninfected mice. FoxP3 staining confirmed over 93% purity of the Tregs (S.Fig. 3A). We exposed *Pink1****^−/−^*** and WT Tregs to the *H. pylori* sonicate and analyzed both the frequency and intensity of FoxP3 by flow cytometry. While frequency of cells expressing FoxP3 dropped overnight to a similar degree in both WT and *Pink1****^−/−^***, the expression level of FoxP3 (Median fluorescence intensity) decreased significantly in *H. pylori* sonicate-treated *Pink1****^−/−^*** Tregs but not in WT Tregs. This suggests that in the absence of PINK1, Tregs have increased susceptibility to FoxP3 loss, both spontaneously and even more in response to *H. pylori* components (Fig. 4B).

Since FoxP3 stability is indispensable for Treg function, the observed phenotypes of the *Pink1****^−/−^*** Tregs suggested they might be less functional in T cell suppression. To address this question, we isolated splenic Tregs from *Pink1****^−/−^*** and WT mice and added them to the 2C TCR CD8 T cell MitAP assay described in Figure 1 (Fig. 4C). Whereas CD25+FoxP3 splenic Tregs from WT mice were effective in suppressing 2C CD8 T cell proliferation induced by *H. pylori-* stimulated BMDC of both *Pink1****^−/−^*** and WT mice, the Tregs from *Pink1****^−/−^*** mice were significantly less suppressive and even slightly promoted 2C CD8 T cell proliferation in some conditions (Fig. 4D-E). We also assessed the capacity of Tregs from *Pink1****^−/−^*** and WT mice to suppress CD4 T cell responses using SMARTA TCR transgenic mice that encode a CD4 T cell receptor directed against a specific LCMV gp61-80 peptide (Oxenius et al., 1998). Indeed, similar to what was seen in the CD8 suppression assay, Tregs from *Pink1****^−/−^*** mice were less potent in suppressing peptide-activated CD4 T cell proliferation than WT Tregs (S.Fig. 3B-C).

Together, these results demonstrate that Treg cells in mice deficient in PINK1 are less effective than WT Tregs at suppressing autoreactive, mitochondria specific CD8 T cells, and that infection with *H.* pylori directly affecting FoxP3 expression in PINK1 deficient Tregs might further compromise Treg function.

### The H. pylori-induced behavioral phenotypes in PINK1 KO mice are CD8 T cell-dependent

Having uncovered two major immunological phenomena developing in response to *H. pylori* infection in *Pink1****^−/−^*** mice, namely an increase in the number of autoreactive CD8 T cells and a loss of T regulatory cell suppressive function, we next investigated whether CD8 T cells could be the main driver associated with the motor dysfunction. For this purpose, we depleted peripheral CD8 T cells in *Pink1****^−/−^*** and WT mice prior to infection with *H. pylori* (Fig. 5A). Mice received three intraperitoneal injections of anti-mouse CD8a and depletion of the CD8 T cells was confirmed by staining of murine blood 24 hours after the last injection and 24 hours prior to infection (Fig. 5B). By two months of infection, CD8 T cell in periphery started to replenish and did so equally in mice of both PINK1 genotypes as seen by the proportion of CD8+ T cell in the spleen (Fig. 5B).

**Figure 5.**
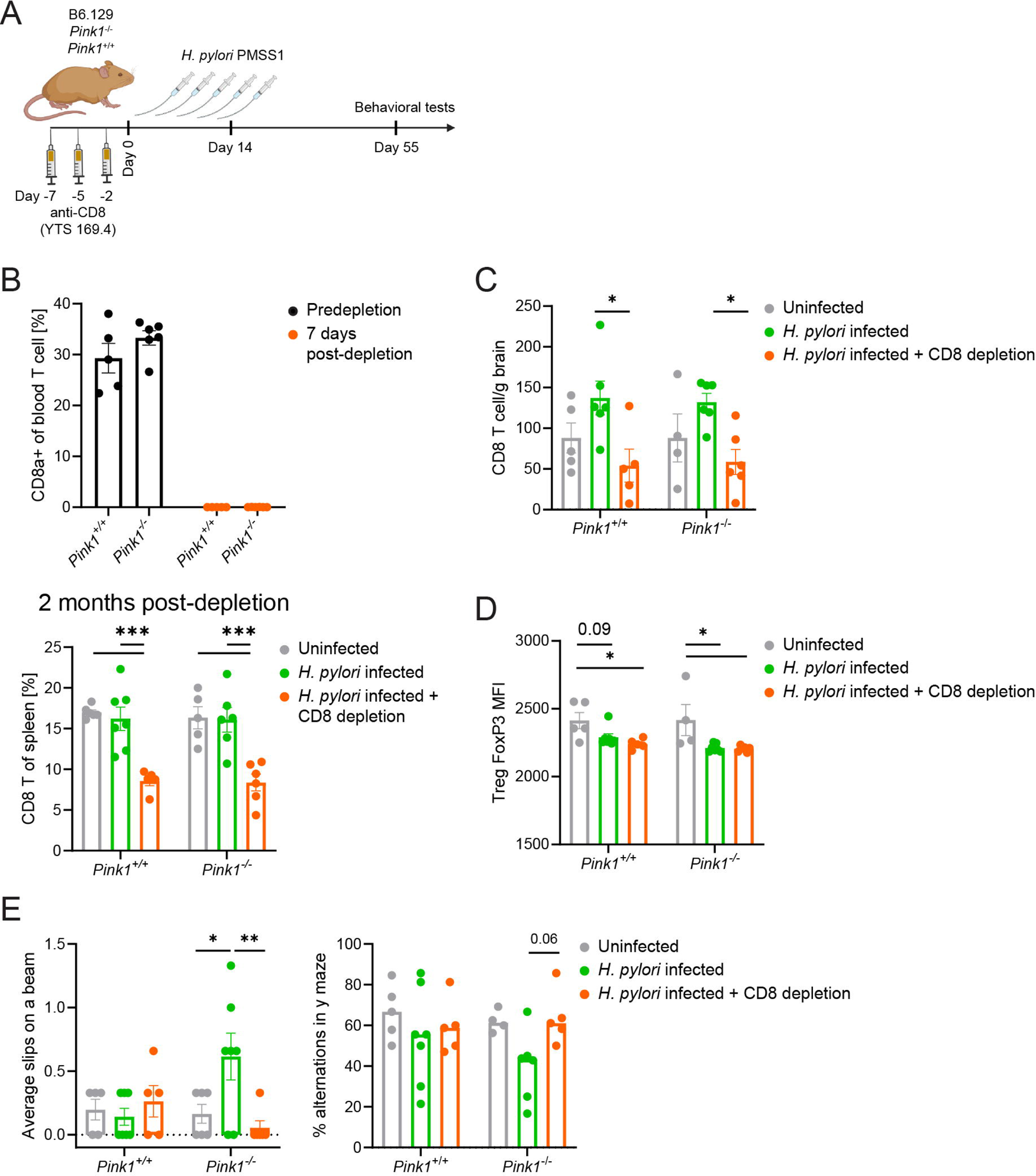
CD8 T depletion in *Pink1^−/−^* mice ameliorates motor phenotype after *H. pylori* infection. A. Experiment design. Six to 10-week-old *Pink1^−/−^* or WT (*Pink1^+/+^*) littermate mice had their peripheral CD8 T cells depleted by intraperitoneal injections of 250 ug of anti-CD8a - YTS 169.4 every 2-3 days for a total of three injections, or alternatively received PBS mock injections. Two days after the third injection, mice were infected with *H. pylori* PMSS1 strain by oral gavage in a regimen of 5 gavages within 2 weeks. After 2 months of the first gavage, mice were tested for motor-behavioral tests and then sacrificed and had their brains and spleens harvested for further evaluations. B. Top -Frequency of CD8 T cell of blood total T cells pre- and post-depletion. Bottom - Frequency of CD8 T cells in spleen at 2 months post-infection. C. Before sacrifice, mice were injected with anti-CD45 in PBS. Brains were collected and immune cells were isolated, stained for flow cytometry, and brain infiltrating CD8 T cells were accessed. D. MFI of FoxP3 in T reg cells. E. Motor functional and behavioral tests. Left - average number of slips on a beam out of three attempts in a beam test; Right - Y-maze - percentage of alternations done in 5 minutes. Data are represented as mean ± SEM and analyzed by two-way ANOVA with Sidak post-test. *p<0.05, **p<0.01 or exact p value indicated in the figure. n=4-6 mice per group. One depletion experiment is finalized by the time of submission.

As seen previously (Fig. 2C-D), the infected *Pink1****^−/−^*** mice that did not have their CD8 T cells depleted prior to infection developed behavioral signs at 2 months post-infection: in this case a significant increase in slips when crossing the beam and a tendency to have decreased alternations in the Y maze (Fig. 5C). Strikingly, pre-infection depletion of CD8 T cells prevented the development of phenotypes: both significant reduction in slips and a tendency to %alternations in Y maze were observed (Fig. 5E). Neither infection nor CD8 T cell depletion had any impact on these phenotypes in WT mice. The number of CD8 T cells infiltrating the brain increased similarly in non-CD8-depleted mice of both genotypes at two months post-infection, whereas mice that had undergone CD8 T cell depletion prior to infection displayed a similar levels of the brain-infiltrating CD8 T cells as an uninfected mice (Fig. 5C).

Analysis of the Tregs in the infected non-depleted or CD8-depleted mice revealed decreased expression of FoxP3 in Treg cells following *H. pylori* infection regardless of whether the CD8 T cells were depleted or not (Fig. 5D). This suggests that Treg dysfunction on its own is not a self-sufficient driver of the motor phenotype of *H. pylori* infected *Pink1****^−/−^*** mice but does not preclude a role for Treg dysfunction in contributing to motor phenotype development, potentially through its diminished inhibitory effects on CD8 T cells. In contrast, the loss of motor phenotype development upon depletion of CD8 T cells suggests that CD8 T cells play a major role in pathophysiology resulting in the early behavior phenotypes following infection.

## Discussion

We have previously shown that recurrent infection with the mouse specific pathogen *C. rodentium* promotes development of a PD-like motor phenotype in otherwise asymptomatic mice lacking the PD-associated gene *Pink1* (Matheoud et al., 2019). In this study, we developed a new model of induction of a prodromal PD-like phenotype in these mice using the PD-relevant human pathogen *H. pylori*. Another group has previously shown *Helicobacter* family mouse-specific bacterium – *Helicobacter hepaticus-* to aggravate symptoms in spontaneously developing PD-like motor phenotype and synucleonopathy in α-synuclein over-expressing mice (Ahn et al., 2023). Here we show for the first time the relevance of the human PD-associated pathogen *H. pylori* in the development of motor and behavioral symptoms in mice carrying a mutation in a PD-associated gene *Pink1*. Notably, uninfected mice deficient in PD associated genes like *Pink1* and *Parkin,* do not develop motor dysfunction, indicating a requirement for the interaction between genetic susceptibility and an environmental trigger for the development of disease signs and symptoms.

Several mechanisms may explain why *H. pylori* infection is a risk factor for PD. *H. pylori* neutralizes stomach acid, which can lead to intestinal microbiota changes, and the intestinal microbiome has been implicated in PD (Liu et al., 2003; McGee et al., 2018; Saha et al., 2010; Wei et al., 2024). *H. pylori* can mediate chronic inflammation in certain individuals, and inflammation is increasingly implicated in PD (Dutta et al., 2008; Wei et al., 2024). *H. pylori* colonization has also been documented to alter the bioavailability of L-DOPA, a dopamine precursor commonly used to treat the dopamine deficiency in PD patients, suggesting it can also impact disease treatment (Wei et al., 2024). Here we show that *H. pylori* induces MitAP and the induction of autoreactive CD8 T cell responses. Notably, MitAP was previously confirmed to be driven by LPS activation of Toll-like receptor 4 (TLR4) on antigen presenting cells. *H. pylori* is known to have a series of modifications to its LPS lipid A core, which results in reduced TLR4 interaction, leading to it signaling primarily through TLR2 (Varga & Peek, 2017). Further investigation is needed to understand whether the induction of MitAP by *H. pylori* requires TLR4 or involves an independent initiating signaling event.

Over the two months that we assessed these mice following infection, we detected relatively subtle but significant phenotypic changes in the infected *Pink1****^−/−^*** mice: mainly slips and missteps while crossing a horizontal beam and a decreased propensity to systematically explore new environments. PD is known to be a progressive disease, with overt motor phenotypes developing only when most of dopaminergic neurons in the *substantia nigra* are already lost. Decades prior to diagnosis, people that will eventually be diagnosed with PD gradually develop a series of prodromal signs and symptoms, including constipation, behavior changes, depression and an increased frequency of bone fractures that is independent of any detectable changes in bone structure. We hypothesize that the increase in bone fractures in prodromal PD is related to subtle changes in motor coordination, and that this is modeled in our mice in the beam crossing test. We also speculate that if followed for longer time points after infection, the phenotype of *H. pylori*-infected mice may progress to include more overt motor signs.

In the current study, we demonstrate that *H. pylori* can induce self-mitochondrial antigen presentation by dendritic cells, both *in vivo* and *in vitro*, and that this process is regulated by PINK1. *Pink1****^−/−^*** mice infected with *H. pylori* developed higher amounts of mitochondria-reactive CD8 T cells and *Pink1****^−/−^*** BMDCs exposed to *H. pylori* induced strong proliferation and activation of primary mitochondria-reactive CD8 T cells. Moreover, *H. pylori* infection in mice induced a dysfunctional phenotype in regulatory T cells (Beigier-Bompadre et al., 2011). Previously PINK1 deficiency in *in vitro* generated induced Treg (iTregs) has been linked to a less suppressive phenotype with lower CD25 and CTLA-4 expression (Ellis et al., 2013). Here, we show for the first time that in the absence of PINK1, ex-vivo Tregs that have developed naturally in mice show a loss of suppressor function towards both conventional CD4 and autoreactive CD8 T cells, primed by *H. pylori* exposed BMDCs.

Even though they were subtle, the motor-behavioral phenotypes in *H. pylori*-infected *Pink1****^−/−^*** mice highly correlated with the presence of autoreactive CD8 T cells, Treg dysfunctional phenotype and brain infiltration with CD8 T cells, suggesting that this complex shift toward immune autoreactivity is associated with disease development. We demonstrated that CD8 depletion prevented the development of the motor behavioral phenotype in *Pink1****^−/−^*** mice, while the Treg dysfunctional phenotype was still induced by the infection regardless of CD8 depletion. This observation reveals a dominant role of the CD8 T cells in the motor behavioral changes, though the details of the mechanisms by which CD8 cells drive these changes remain to be defined. We hypothesize that Tregs are likely to amplify the overall immune dysregulation in *H. pylori*-infected *Pink1****^−/−^*** mice by their impaired suppression of the autoreactive CD8 T cells, thus exacerbating disease development. The role of Tregs in PD has been under focus recently and several clinical trials targeting the Treg population have been initiated (Olson et al., 2023; Park et al., 2023). The mouse model described here provides a relevant preclinical system to assess the effects of Treg modulation in future studies.

Of note, we did observe some variability in certain parameters between mice and between experiments. We hypothesize that the variance in mouse performance and in some immunological parameters like baseline brain infiltration may be due to the fact that these mice are on a mixed genetic background and may also have microbiome differences, both of which are known to impact PD. Indeed, despite the increased variance, this may better represent the genetic and microbiome diversity in human populations and favor the detection of the most robust effects.

Overall, we show that exposure of PD-susceptible animals to the PD-associated microbe *H. pylori* leads to the development of significant motor and behavioral phenotypes that resemble those seen in prodromal PD and are dependent on CD8 T cells. Though many questions remain, we anticipate that a model integrating genetic PD susceptibility, a PD-relevant environmental trigger, and specific immune changes that are required for symptom development will be a valuable tool for increasing our understanding of this complex disease.

## Methods

### Bacteria

*H. pylori* PMSS1 strain was a kind gift from Dr.Russell Jones. Bacteria were cultured on Columbia agar supplemented with 5% horse blood, Vancomycin, Cefsulodin, Polymixin B, Cycloheximide, Trimethoprim, Amphotericin B, and β-cyclodextrin. To prepare the inoculum, bacteria were grown in Brain Heart Infusion broth supplemented with 10% horse serum and 0.25% yeast extract for 48 hours with shaking at 37 degrees C in T75 tissue flasks with venting cap in a standing position. Microaerophilic conditions were generated with BD Campy Gas Packs.

### Mice

All animal experiments were performed under conditions specified by the Canadian Council on Animal Care and were approved by the McGill University Animal Care Committee. *Pink1* heterozygous mice on a mixed B6.129 background were bred to generate knockout (*Pink1−/−*) and wild-type littermate control mice, the latter were used in experiments as previously described (Matheoud et al., 2019).

2C TCR mice on a C57BL/6 background were donated by Dr. Nathalie Labrecque. OT-1 TCR TgXRag1−/− and SMARTA-1 TCR Tg mice were a gift from Dr. Heather Melichar.

### Infection

Six to 10-week-old mice were starved for 3 hours prior to gavage with free access to water. Mice were given 0.1 ml of 0.1 M sodium bicarbonate by oral gavage, followed with either 0.1 ml of sterile Brain Heart Infusion (BHI) broth for the group of uninfected mice or with 0.1 mL of BHI broth containing 1×10^9 CFU *H. pylori* PMSS1 strain five times within two-week.

### In vivo CD8 depletion

For depletion of peripheral CD8 T cell PINK1^−/−^ and WT were randomized into two sex and age equivalent groups, CD8 depletion group received three i.p. injections on 7, 5 and 2 days prior to *H. pylori* infection (250ug/mouse in 200 ul of PBS; clone YTS169. Purified in vivo GOLD™, Leinco Cat#C2442). The comparison group received three i.p. injections of PBS. Efficiency of CD8 depletion was confirmed with flow cytometry of blood obtained by a cheek vein bleeding on day 8 and day 1 prior to infection. The mice of the comparison group were also bled without further processing of the blood.

### Quantitative PCR for *H. pylori*

To assess bacterial colonization, half of the antrum and corpus portion of the stomach were homogenized in BHI. Genomic DNA was extracted using the GeneJet Genomic DNA purification kit according to manufacturer’s instructions (Thermo Scientific, Waltham, Mass). qPCR was performed on StepOnePlus (Applied Biosystems, USA) using two primers and an internal probe specific to *H. pylori UreaseB (UreB)* as previously described (Fox et al., 2007). A standard curve was generated using serial 10-fold dilutions of *H. pylori* PMSS1 genome copies using an estimated mass of 1.66Mb. UreB levels were normalized to murine DNA as quantified by a qPCR assay for 18s rRNA (forward, 5′- GCCTGCGGCTTAATTTGAC -3′, and reverse, 5′- CGGAATCGAGAAAGAGCTATCA -3′), and an internal probe (5′- CGGACACGGACAGGATTGACAGAT -3′).

### Histology

To examine stomach histology, a portion of the antrum and corpus of the stomach was fixed in formalin and embedded in paraffin. 4-micron sections were prepared, deparaffinized, and hydrated followed by hematoxylin and eosin staining.

### Cell culture

2cz CD8 T Hybridoma cells were cultured in complete RPMI media (Gibco): media supplemented with 10% FBS (Wisent, Cat#090150), and Wisent’s 10mM Hepes, 2mM L-glutamine, 1mM Sodium Pyruvate and 100 IU Streptomycin/Penicillin.

Bone marrow derived dendritic cells (BMDC) were generated from the femur’s bone marrow single cell suspension,n cultured in the complete RPMI with 20 ng/ml of rmGM-CSF (Biolegend, cat#576308) for 9 days.

5×10^6 Murine splenocytes after ammonium chloride potassium buffered Red Blood Cell lysis were incubated with/without *H. pylori* sonicate 20-80 ug/ml. After 24 hours cells were pelleted by centrifugation, supernatant collected for ELISA and cells were used for Flow cytometry.

### Mitochondrial Antigen Presentation (MitAP) to 2cz hybrydoma

MitAP was assessed by flow cytometry of the activation induced marker (PD1/CD69/CD137 - AIM) expression on 2cz hybridoma cells after co-culture with BMDC. In brief, 2cz were stained with 1uM Tag-it Violet (Biolegend, Cat#455101) in PBS for 12 minutes at 37C, blocked with FBS 1:1 for extra 5 minutes at RT and washed three times in complete RPMI to obtain 2cz-Tag-it. 2cz-Tag-it alone served a negative control. 2cz-Tag-it co-culture with RAW 264.7 cells activated for 6 hours with 1.5 ug/ml Ultrapure LPS from *E.coli* O111:B4 (InvivoGen, Cat#tlrl-3pepls) and 2cz cultured with 50 ng/ml of its strong agonist peptide SIYRYYGL (GenScript) were used as a positive control (Matheoud et al., 2019; Udaka et al., 1996). BMDC of three *Pink1^−/−^* and three littermate WT mice were treated for 6 hours with 1.5 ug/ml LPS as above, recombinant *H. pylori,* CagA 1 ug/ml (my Bioscourse, Cat#MBS596143), live *H. pylori,* or heat killed for 30 minutes at 60C *H. pylori* (HK *H. pylori*) at MOI of 100 or sonicate of *H. pylori* 100 ug/ml.

2cz cells were cultured with *H. pylori* exposed or untreated BMDC for 24 hours at 1:2 starting ratio, afterwards cells were collected and used for Flow cytometry.

The average value of AIM expression of technical triplicates for each condition was used for analysis.

### MitAP to primary 2C CD8 T (proliferation and suppression) assay

Spleens from 2C TCR transgenic mice of 6-8 weeks old were mashed through a 70 um cell strainer. RBC lysed as described above. CD8 T cells isolated with a negative magnetic selection using EasySep™ Mouse CD8+ T Cell Isolation Kit (Stemcell) according to the manufacturer protocol. 2C CD8 T cells were stained with 2.5 uM cell tracker Tag-it Violet (Biolegend) in the same manner as 2cz hybridoma.

BMDC were generated as previously described individually from three *Pink1^+/+^* and three *Pink1^−/−^* mice. On day 9 of mouse recombinant GM-CSF culture, BMDC were collected and split in half, for untreated and *H. pylori* sonicate 100 ug/ml exposure. After six hours BMDC were collected with cell scraper and washed in an ice-cold PBS. Washed BMDC were fixed with 1% PFA for 9 minutes at the room temperature, followed by three wash steps in 0.1M Glycine in complete RPMI.

CD4CD25-(CD4 conv) and CD4CD25 T (Tregs) cells were isolated from spleens of three *Pink1^+/+^* and three *Pink1^−/−^* mice using CD4+CD25+ Regulatory T Cell Isolation Kit (Miltenyi Biotech) according to manufacturer’s protocol. CD4 conv T cells were resuspended in complete RPMI 10% FBS and incubated for 1 hour at 37C with 1 ug/ml of anti-mouse CD3 (2c11 clone; Biolegend) or left untreated. Cells were washed in PBS and stained with CFSE 1.25 uM in the same manner as Tag-it violet staining.

Purity of magnetic selection of 2C CD8, CD4+CD25+ and CD4+CD25-cells was determined with flow cytometry prior to co-culture. Regulatory CD25+CD4 T cells were stained with anti-FoxP3 and acquired with flow cytometry.

Co-culture was performed in 10% FBS complete RPMI in 96-well round bottom plates for suspension cells (Sarstedt) at a following ratio: 50,000 2C to 50,000 CD4 conv cells to 100,000 fixed BMDC. For suppression assay Tregs were added to the co-culture at (1-0.25):1 2C CD8 T ratio. Each condition was performed in technical replicates. At 24- and 72-hours cells were collected and assessed by flow cytometry for AIM expression and proliferation.

Suppression of proliferation was calculated using the following formula:

% suppression = 100x(1-proliferation of 2C in the presence of Treg/ proliferation at No Treg condition)

### CD4 T cell proliferation and suppression assay

CD4 conv were isolated from the spleens of SMARTA TCR Tg mice (a generous gift from Dr. Heather Melichar, McGill) using the EasySep™ Mouse CD4+ T Cell Isolation Kit (Stemcell). Cells were washed in PBS and stained with 2.5 Tag-it Violet cell tracker (Biolegend) as previously described.

BMDC of *Pink1* WT mice were generated as previously described and maturated with 1.5 ug/ml Ultrapure LPS from *E. coli* O111:B4 (InvivoGen, Cat#tlrl-3pepls) for six hours. BMDC were washed from LPS and added to the co-culture with the LCMV gp61-80 peptide (Anaspec) to a final concentration in co-culture of 2.5 ug/ml.

CD4CD25+ T (Tregs) cells were isolated from spleens of three *Pink1^+/+^* and three *Pink1^−/−^* mice using CD4+CD25+ Regulatory T Cell Isolation Kit (Miltenyi Biotech) according to manufacturer’s protocol.

Purity of magnetic selection of SMARTA CD4 conv, CD4 CD25+ was determined with flow cytometry prior to co-culture. CD25+ cells were also stained for FoxP3 and acquired with flow cytometry.

Co-culture was performed in 10% FBS complete RPMI 96-well round bottom plates for suspension cells (Sarstedt) at a following ratio: 50,000 SMARTA CD4 T to 10,000 BMDC and a total volume of 200 ul per well. For suppression assay Tregs were added to the co-culture at 0.5-0.125 to 1 SMARTA CD4 T ratio. Each condition was performed in technical replicates. At 72 hours cells were collected and assessed by flow cytometry for Tag-It Violet signal dilution (indicative of cell division).

Suppression of proliferation was calculated using the following formula:

% suppression = 100x(1-proliferation of SMARTA in the presence of Treg/ proliferation at No Treg condition)

### *Ex vivo* autoreactive mitochondrial antigen (MitAg) specific CD8 T cell assay

To generate antigen-presenting cells (APC) – BMDC of *Pink1^−/−^* mice were stained with Tag-it Violet as described previously and then coated on ice for 3 hours with the mitochondrial peptide pool (custom synthesis by Genscript or Canada Peptide) or mock pool (OVA 257-264, LCMV Gp33-41; OVA 323-339 GenScript) 5 ug/ml each.

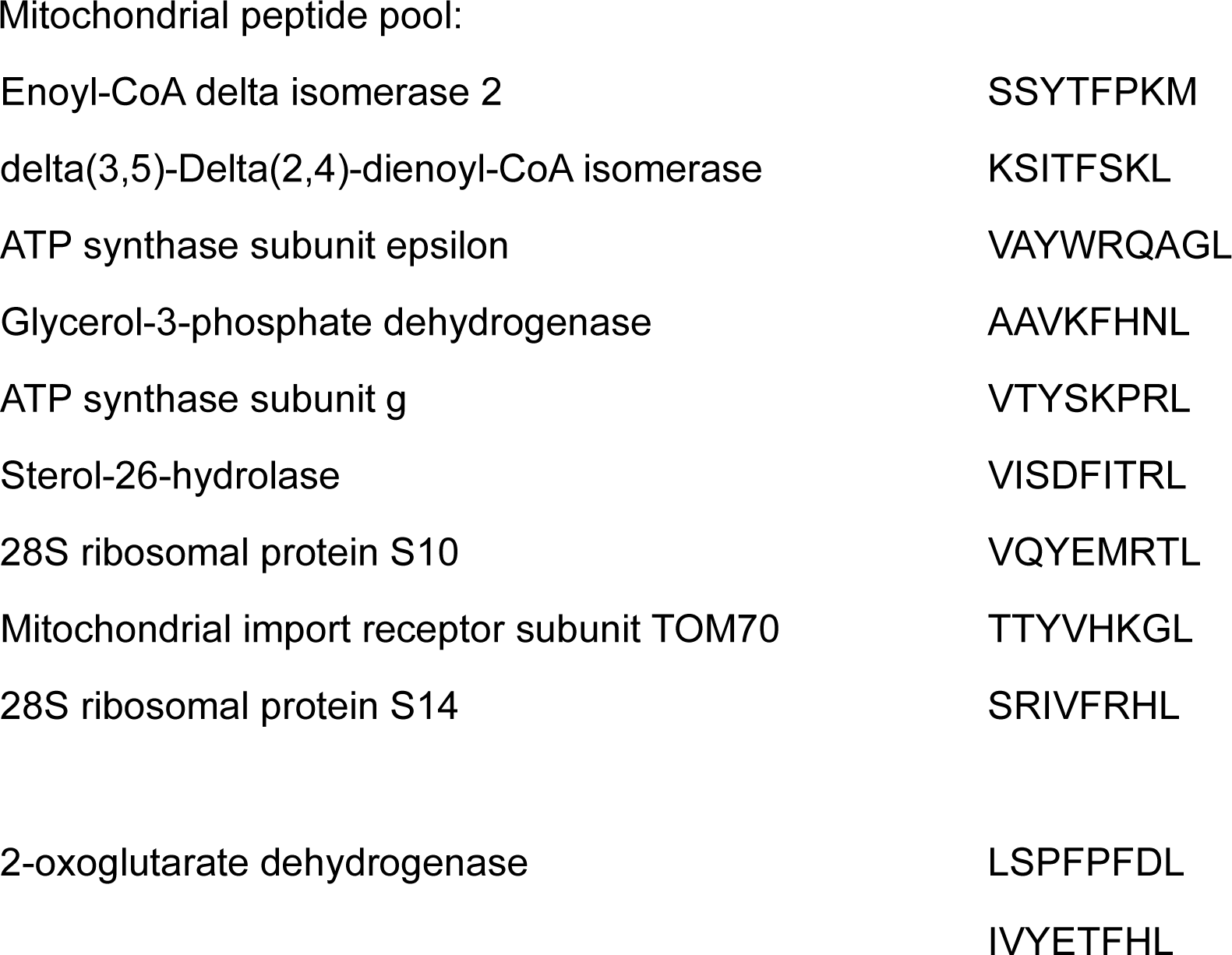
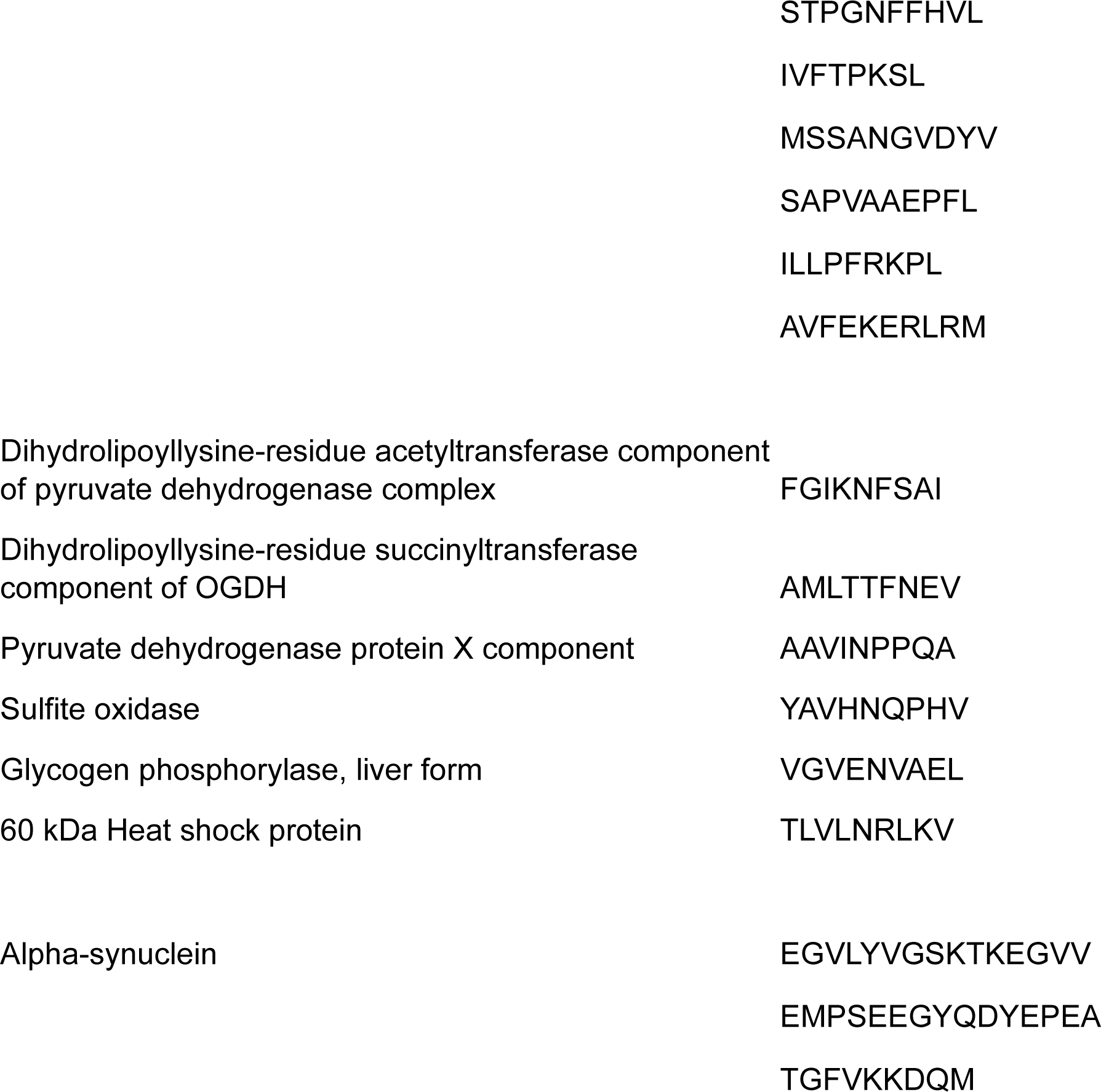

BMDC were washed in ice cold PBS and fixed at RT in 1% paraformaldehyde for 9 minutes, followed by three wash steps in 0.1M Glycine in complete RPMI. Splenocytes of mice were cocultured with APC-MitAg or APC-mock in 5:1 ratio overnight. Cells were collected for flow cytometry. Activation induced marker (PD1/CD69/CD137) expression on TCRb+CD8+ in APC-MitAg condition over APC-mock condition was used to determine frequency of autoreactive MitAg-specific CD8 T cells.

### Flow Cytometry

Cells were stained with 1:2000 in Zombie red fixable viability dye (Biolegend, Cat#423109) 20 min on ice, followed by Fc-Block step with anti-mo CD16/CD32 (Invitrogen, Cat# 14-01061-86) and staining of surface markers with a mixture of conjugated antibodies 1:250 in 2%FBS PBS 1mM EDTA buffer with 10% Brilliant Stain Buffer Plus (BD, Cat#566385) for 30 min on ice. For transcriptional factor (TF) staining cells were fixed in Foxp3/Transcription Factor Staining Buffer (eBiosciences, cat#00-5523-00) for 60 min on ice, then washed in permeabilization buffer and stained with a mixture of conjugated antibodies in the same buffer 1:100-1:250. Single stained cells of the appropriate tissue processed accordingly were used as a reference control for unmixing, for a single stain of TF UltraComp eBeads (Invitrogen, Cat#01-2222-42) were used. For autofluorescence deduction unstained cells of the appropriate tissue were acquired. Samples and controls were acquired using spectral cytometer Aurora 4L (Cytek) and conventional LSRII Fortessa 5L (BD). For spectral cytometry unmixed in SpectroFlo FCS 3.0 files exported and analyzed using FlowJo v10.9 Software (BD).

#### Antibodies

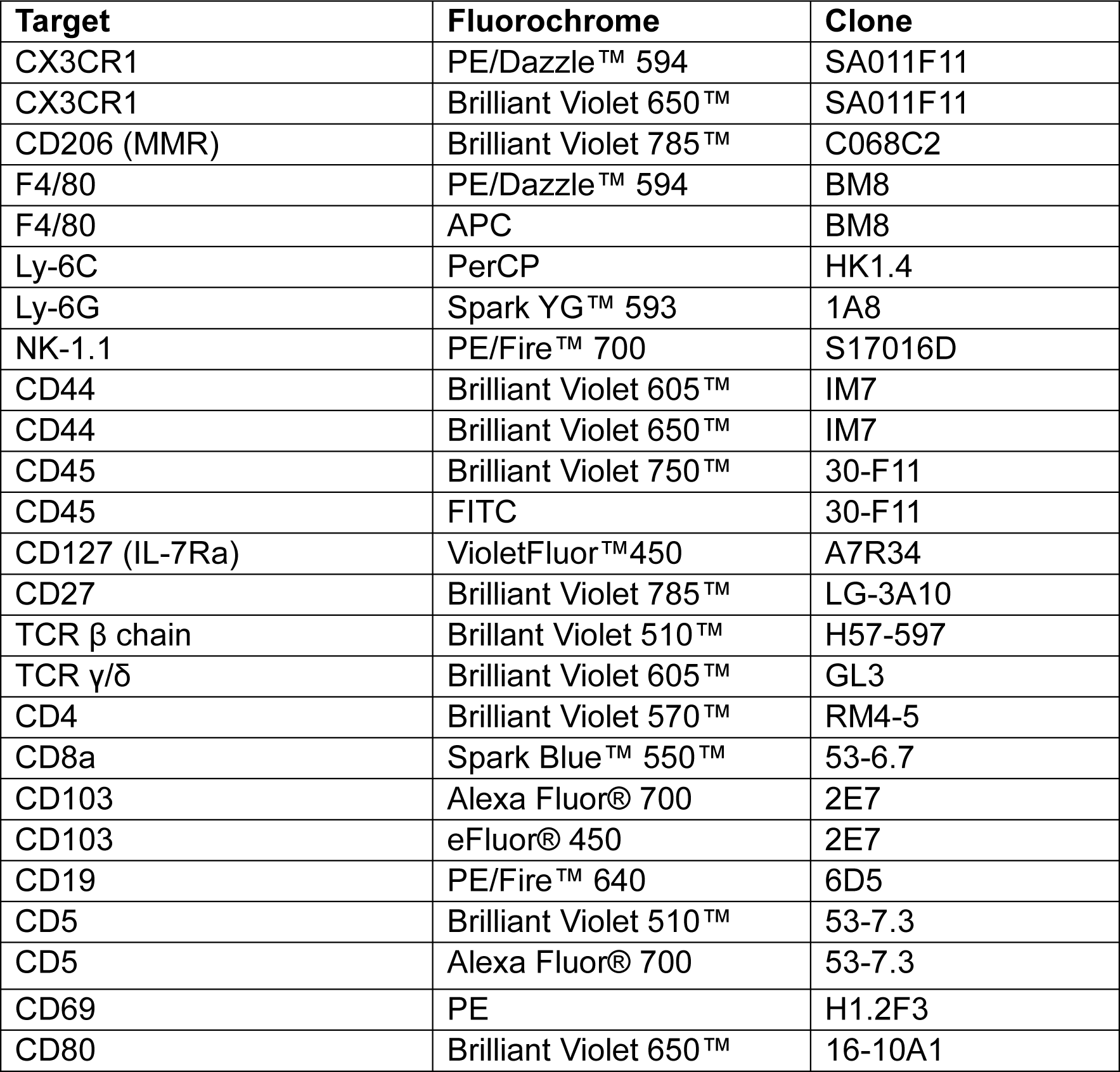
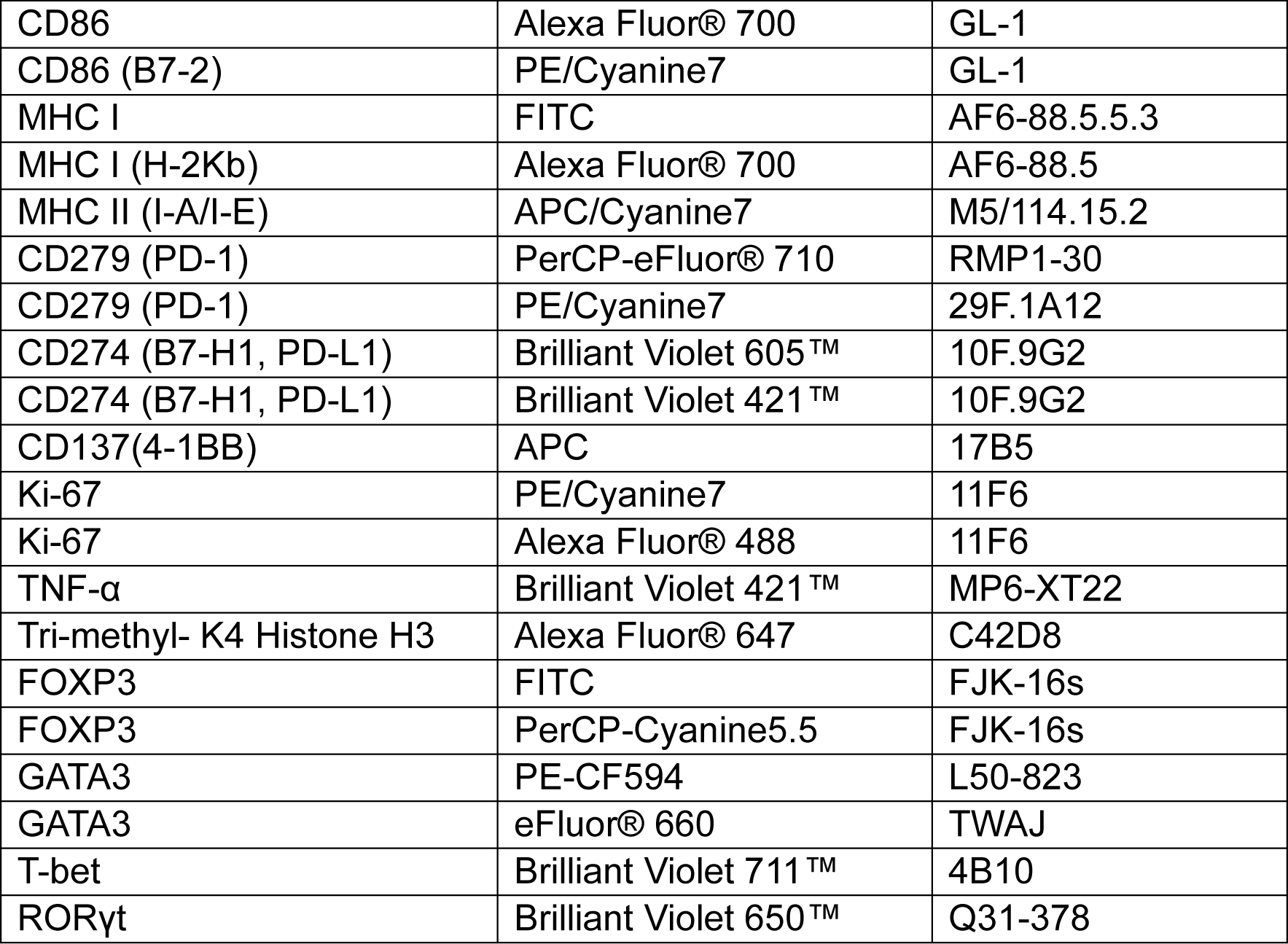

### Brain infiltrating leukocytes (BIL) extraction

For BIL extraction mice were injected with 200 ul of anti-mouse CD45-FITC in PBS 6 ug/mouse i.p. and sacrificed after 20 min (time point and concentration assessed previously experimentally with cheek bleeding of mice, data not shown), brains were collected into pre-weighted 15 ml conical tube containing 3 ml RPMI media. Tubes with brains were weighted for the brain mass calculation.

Brains were minced with scissors and dissociated in digestion buffer (RPMI with TL Liberase (Roche, Cat#5401020001) 2.5 ug/ml, DNAse I (Roche, Cat#04416728001) 10 ug/ml and 1.5 mM Calcium Chloride) for 35 min at 37C with gentle shaking every 10-15 minutes. Digestion enzymes were blocked with 10%FBS 2mM EDTA PBS and tissue was filtered through 70 um cell strainers. Immunocytes were isolated using gradient centrifugation with 37% Percoll (Cytiva, Cat#17089102) in D-PBS (with Calcium and Magnesium) at room temperature for 20 minutes no break. Myelin containing upper layer was removed with a pipette and a cut 1ml tip. Cells in the interphase spin down and the cell pellet was treated with Ammonium Chloride Potassium lysing buffer and processed for flow cytometry.

### ELISA

Tumor necrosis factor alpha in culture supernatants was measured with Mouse TNF-alpha Quantikine ELISA Kit according to manufacturer’s protocol (RnD Systems, Cat#MTA00B).

### Multiplex assay

To assess local immune response, a portion of the antrum and the corpus of a murine stomach was homogenized in B150 lysis buffer followed by sonication three times at 60% amplitude for 8 seconds. The sonicate was then centrifuged and the supernatant was collected for further multiplex analysis. Eotaxin, G-CSF, GM-CSF, IFNγ, IL-1α, IL-1β, IL-2, IL-3, IL-4, IL-5, IL-6, IL-7, IL-9, IL-10, IL-12p40, IL-12p70, IL-13, IL-15, IL-17A, IP-10, KC, LIF, LIX, MCP-1, M-CSF, MIG, MIP-1α, MIP-1β, MIP-2, RANTES, TNFα, VEGF-A in stomach tissue homogenates were assessed using Mouse Cytokine/Chemokine 32-Plex Discovery Assay® Array (MD32) (EveTech). Measurement was performed in duplicates for each sample.

### Mouse Behavioral tests

#### Y-maze

Mice were placed in arm B of the Y maze (Harvard Apparatus Cat#76-0079) and left to explore the maze for 5 minutes. The number of entries in any of three arms (A, B or C) and alternations were calculated. All four limbs are within the arm were considered as an entry. % Alternations were calculated by dividing alternations by the total number of possible triads and multiplied by 100. The number of fecal pellets left in the maze was recorded. The maze was cleaned after each mouse with 0.5% Peroxigard to remove all mouse traces and odor, and for disinfection.

#### Pole test

For the pole test, mice were first transferred to the cage with the pole in it and left to explore for 5 minutes. After acclimatization, the mouse was first put on the top of the pole head downwards and let descend for three trials (descend time was recorded). Then a mouse was placed on the top of the pole head upwards and left to turn and descend. The time taken to fully place the hind legs over the front ones on the pole was recorded as a T-time. An average of three attempts to descend and turn was used for statistical analysis.

#### Beam test

Mice were trained to cross the beam the day before the test, three attempts per mouse. On the trail day the time to cross the one-meter cylindrical beam was recorded, slips of the legs on the beam were also recorder for each attempt. An average of three attempts to traverse the beam, and average number of slips was used for statistical analysis.

### Statistical analysis

Statistical analysis was performed using GraphPad Prism 8.0.1 (Dogmatics). P-value<0.05 was considered a significant difference and marked as *; p<0.01 - ** and p<0.001 as ***. Each experiment was repeated at least two times with a minimum of three mice per group (except CD8 depletion experiment). All *in vitro* assays were performed at least twice, each individual experiment in technical replicates plicates, with at least three biological replicates within each group.

### Illustrations

Graphical abstract and experimental designs illustration were created with BioRender.com.

## Acknowledgements

The study was funded by the joint efforts of The Michael J. Fox Foundation for Parkinson’s Research (MJFF) and the Aligning Science Across Parkinson’s (ASAP) initiative. MJFF administers the grant ASAP 000525 on behalf of ASAP and itself. This study was also funded by a CIHR Operating grant awarded to SG.

We claim equal contribution of all co-authors to this paper. We believe that scientific research is only possible as a teamwork, and no matter how great any individual scientist is, together we can achieve more.

## Supplementary Figures

**S.Figure 1. MitAP assay modifications. PINK1 of BMDC does not affect proliferation of conventional CD4 T cells preactivated with anti-CD3, nonetheless it alters the expression of MHC class I and II, and co-stimulatory molecules in BMDCs after *H. pylori* stimulation.**

A. Dose dependent response of 2cz hybridoma: the frequency of activation induced molecules (CD69/CD137/PD-1) to the SIYRYYGL concentration in the culture. Data are represented as mean of technical triplicates. B. Correlation of MitAP detection with flow cytometry acquisition of activation induced markers (AIM) to IL-2 production in 2cz cells activated with LPS-exposed RAW cells detected by ELISPOT. Data are analysed by Pearson correlation (r). (C-D) BMDCs from *Pink1^−/−^* or WT (*Pink1^+/+^*) littermate mice were differentiated with 20 ng/ml rmGM-CSF for 9 days. Then, cells were stimulated for 6 hours with sonicated *H. pylori* and fixed. Fixed BMDC were analyzed by flow cytometry immediately. The remaining BMDC were co-cultured with splenic 2C CD8 T cells labeled with Tag-it Violet cell tracker and CFSE-labeled CD4 T cells, preactivated with anti-CD3 antibody. Co-culture lasted 72 h, cells harvested and stained for flow cytometry. C. Overlaid histograms of CFSE fluorescence (left) of CD4 T c cells and frequency of divided pre-activated with anti-CD3 CD4 T (right). Data are represented as a mean ± SEM of technical replicates and analyzed by two-way ANOVA with Sidak post-test. Each Cell type was extracted from spleens of 3 individual mice per group, pooled together and co-cultured in technical replicates. Experiments were done twice, independently. D. MHC-I, MHC-II and co-stimulatory CD86 and CD80 molecules MFI of CD11c+CD11b+ fixed BMDC. Data are represented as mean ± SEM and analyzed by two-way ANOVA with Sidak post-test. Cells were extracted from 3 individual mice per group, pooled together and analyzed in staining duplicates. **p<0.01, ***p<0.001. Experiments were done twice, independently. E. Pearson correlation of MHCI expression on *H. pylori* exposed BMDC and AIM on 2cz hybridoma.

**S.Figure 2. *Pink1^−/−^* or WT mice exhibited mild gastric pathology after *H. pylori* infection.**

Six to 10-week-old *Pink1^−/−^* or WT (*Pink1^+/+^*) mice were infected with *H. pylori* PMSS1 strain by oral gavage in a regimen of 5 gavages within 2 weeks. After 2 months of the first gavage, mice were sacrificed and had their stomachs harvested for clinical and inflammatory assessments. A. Stomach clinical assessments - diameter, half of stomach area, and pyloric diameter were assessed with ImageJ analyzed of the stomach photographs. Data are represented as mean ± SEM and analyzed by two-way ANOVA with Tukey post-test. n=5-10 mice per group. Two independent experiments are represented. B. Representative stomachs of infected *Pink1^−/−^* (right) or *Pink1^+/+^* (left) infected mice. Scale bars represent 1 cm. C. H&E-stained sections of the stomachs. D. at 2mpi mice were sacrificed and had their stomachs harvested and homogenized for cytokine and chemokine multiplex evaluation. Heatmap of relative to the average of uninfected *Pink1^+/+^* mouse level of cytokines and chemokines in gastric homogenates of mice. Data are represented on a heat map as median and analyzed by two-way ANOVA of all groups against WT group. Significant difference with p<0.05 is indicated in bold cytokine name with a superscripted significantly different group (a - *Pink1^+/+^* infected; b – *Pink1^−/−^* infected mice). The number of mice per group is indicated in the figure. Cytokines were assessed in one experiment.

**S.Figure 3. T reg cells of *Pink1^−/−^* mice exposed to *H. pylori* sonicate lose their suppressor phenotype.**

Six to 10-week-old *Pink1^−/−^* or WT (*Pink1^+/+^*) were sacrificed and spleens were harvested, and flow cytometry of magnetically isolated CD25+ CD4 T cells was performed. A Purity of CD25+CD4 T isolated cells. B. Overlaid histogram of CD4 T cell of SMARTA Tg mice proliferation after cells were activated with agonistic LCMV gp61-80 peptide for 72 h in the presence of BMDC. C Frequency of suppression of CD4 T cells proliferation by Treg cells in the different ratios. Data are represented as mean ± SEM and analyzed by two-way ANOVA with Sidak post-test. Tregs were extracted from three mice per group *p<0.05, **p<0.01. One representative of three independent experiments is showed.

## Notes

### Competing Interest Statement

The authors have declared no competing interest.

