## Supplementary figures and images for "Modeling gene-environment interactions in Parkinson’s Disease: *Helicobacter pylori* infection of *Pink1^−/−^* mice induces CD8 T cell-dependent motor and cognitive dysfunction"

### Supplemental Figure 1

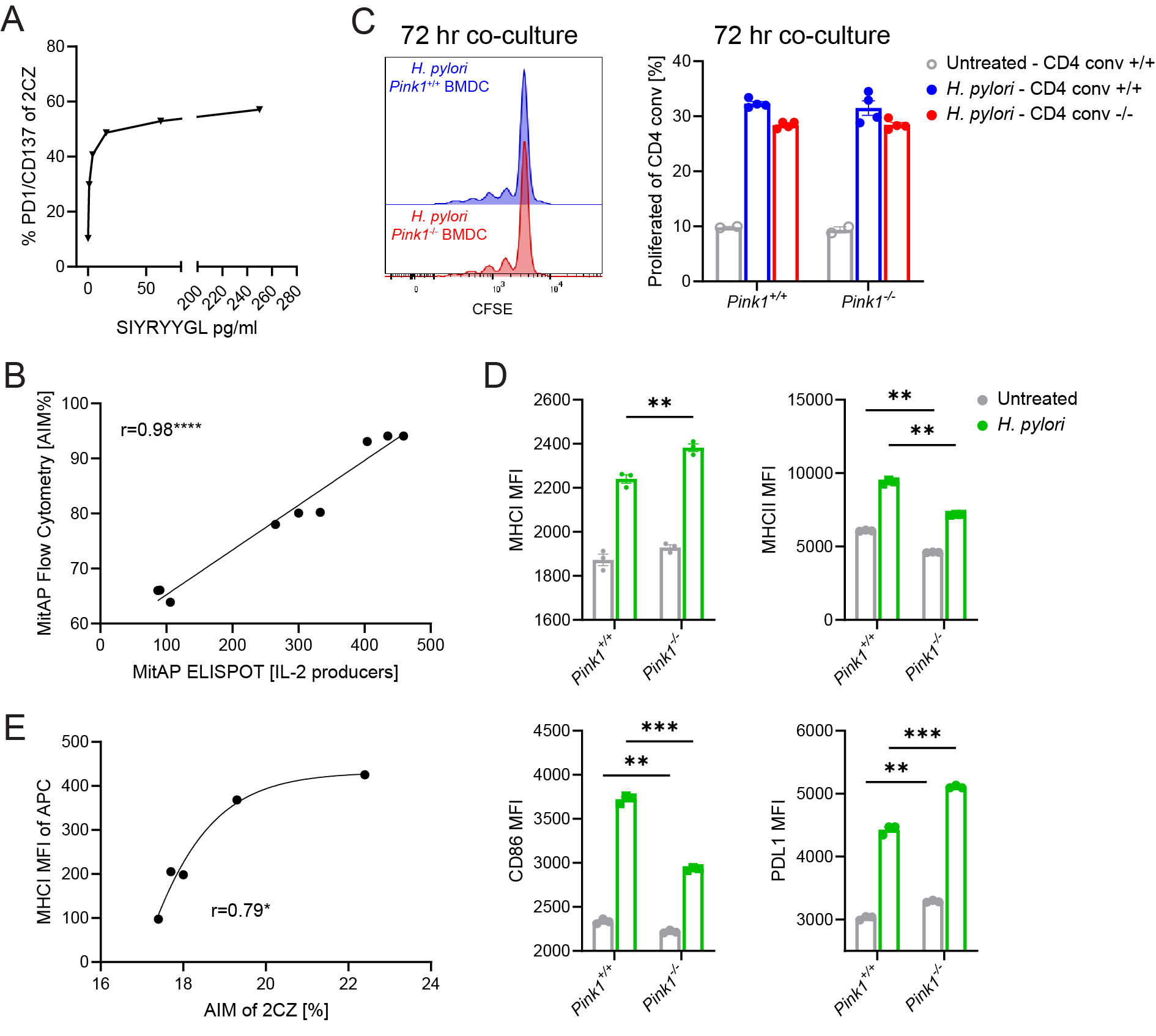

### Supplemental Figure 2

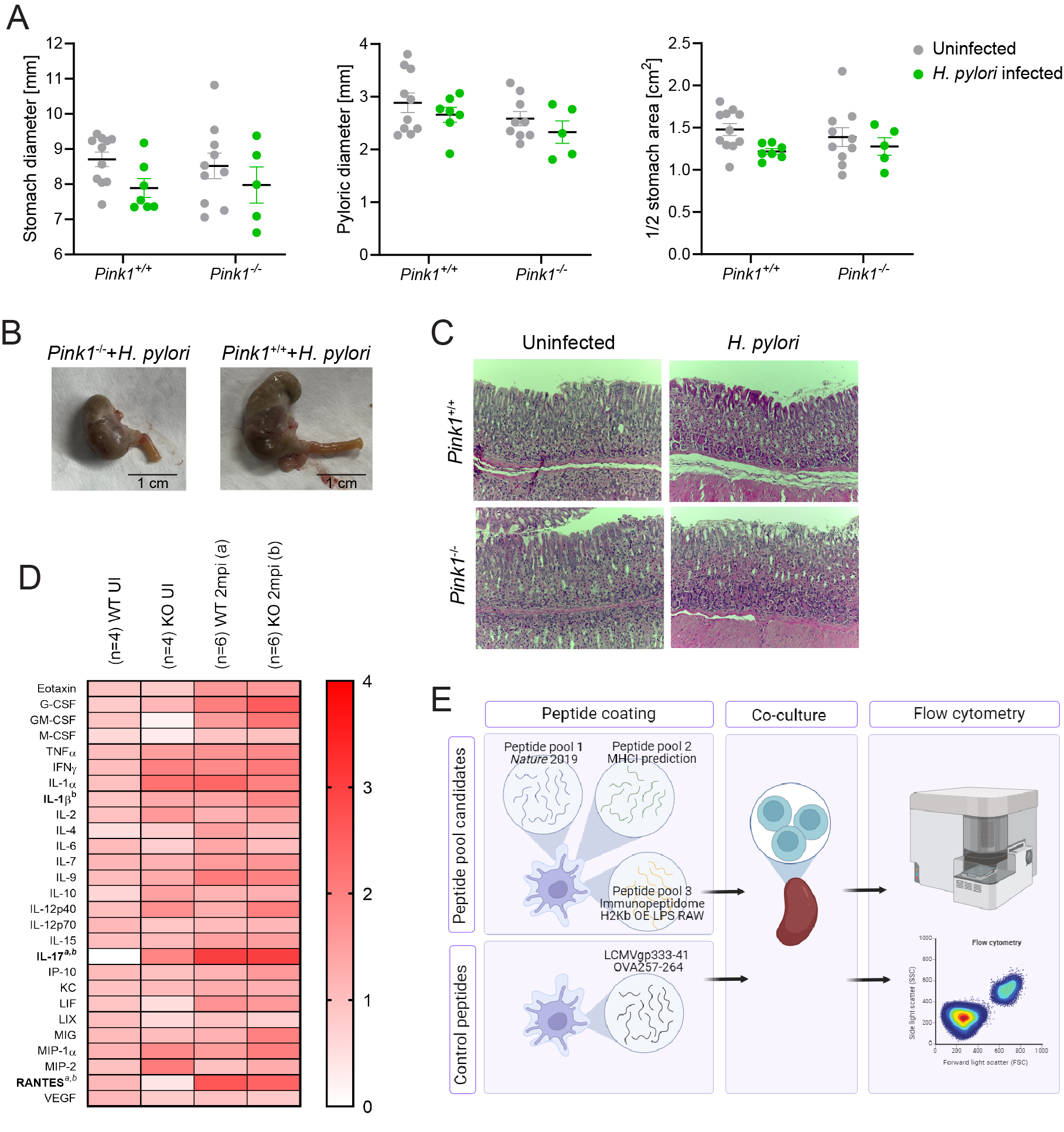

### Supplemental Figure 3

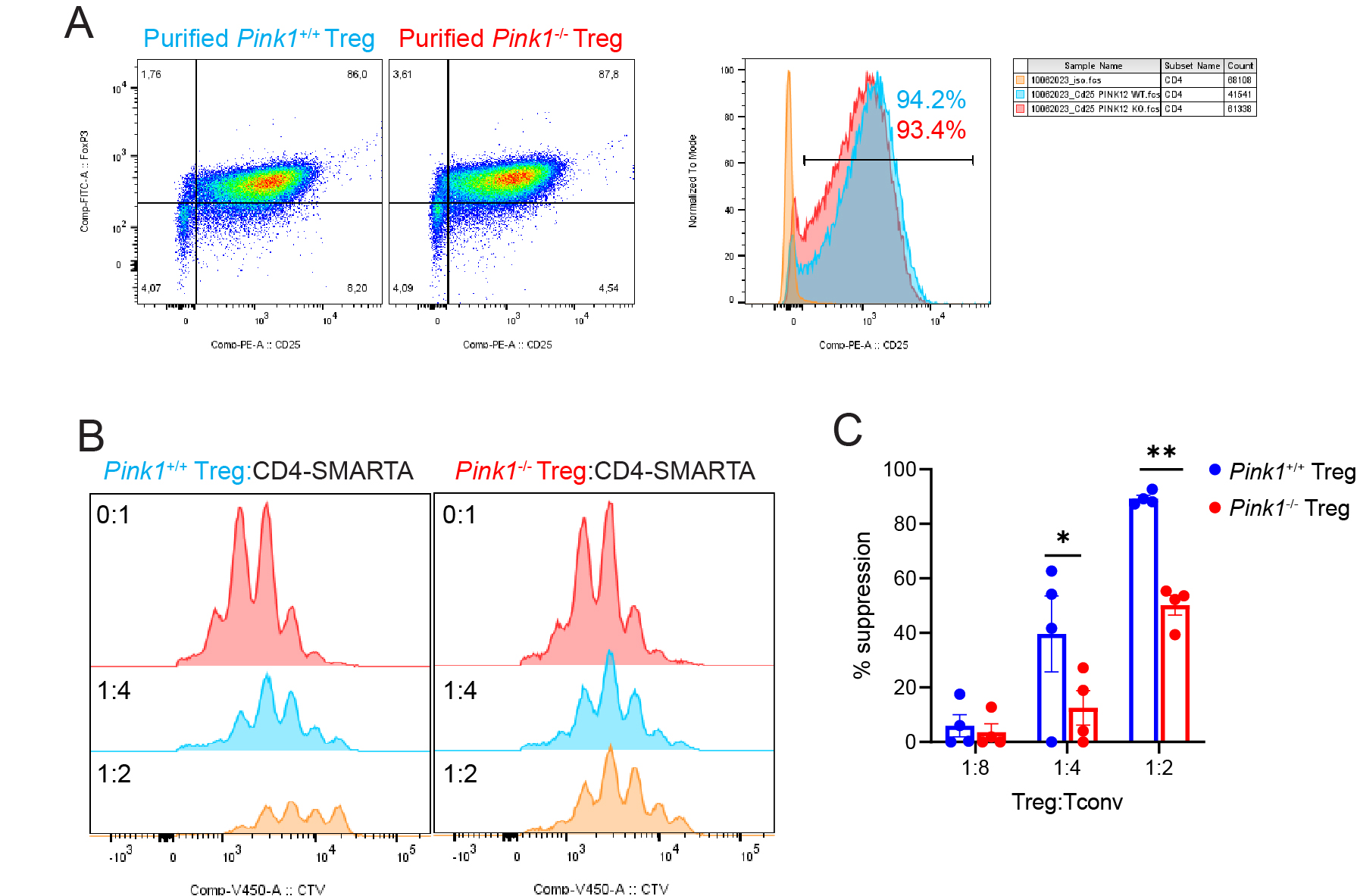
